# Senescence-induced Spatio-temporal Re-organization of the Nuclear Lamina Dynamic Landscape at Single-molecule Level in Live Human RPE Cells

**DOI:** 10.64898/2026.10.08.756399

**Authors:** Xuan Kuo, Wilfried Engl, Pakorn Kanchanawong, Ziqing Winston Zhao

## Abstract

A defining hallmark of cellular senescence is the extensive remodeling of the nuclear lamina (NL) in both structure, composition, and its interactions with lamina-associated genomic domains. However, how such senescence-induced changes could quantitatively impact the spatio-temporal dynamics of NL interactions at the single-molecule level remain poorly understood. Herein, using human retinal pigment epithelium (RPE) cells as model system, we establish a robust, chemically inducible cellular senescence model capable of recapitulating key signatures of the senescence-associated phenotype. Senescent RPE cells exhibit distinct alteration patterns in the expression of various NL components, including lamin B receptor (LBR) and emerin, two key mediators of NL– chromatin interactions. Importantly, live-cell diffusion measurements via imaging FCS reveal globally enhanced mobility for both LBR and emerin, indicative of weakened NL anchoring in senescent cells. Single-molecule tracking and super-resolved STAR mapping further unveil distinct, multi-modal re-organization of their NL-binding dynamics in both space and time (especially in the form of LBR binding “hotspots” where multiple stable binding events cluster to effect more extended and prolonged interaction), in line with their respective expression-level alterations and differential lamin interactions. Moreover, correlative mapping uncovers a strong spatial coupling between reduced LBR mobility and highly condensed constitutive heterochromatin domains at the NL, which become dispersed during senescence. Collectively, these findings demonstrate the re-organization of NL dynamic landscape at single-molecule level as a critical and previously under-explored facet of senescence-induced NL remodeling, and point to a synergistic interplay between physical de-anchoring, dynamic attenuation of stable NL-binding and spatial restructuring of chromatin architecture in modulating NL–heterochromatin contacts to potentially effect global changes in senescence-associated gene expression.

**STATEMENT OF SIGNIFICANCE:** In this study, we combine a suite of quantitative imaging and transcriptomic profiling approaches with complementary capabilities and perform, to our knowledge, the first rigorous characterization of the impact of cellular senescence on the spatio-temporal dynamics of NL interactions at the single-molecule level in live human cells. Our findings not only reveal a key and previously overlooked dimension of NL remodeling during cellular senescence, but also shed first-of-its-kind quantitative insights into a mechanism by which the multi-modal binding landscape at the NL work in concert with higher-order chromatin architecture and NL–chromatin anchoring to modulate the contacts between NL and heterochromatin, which can in turn alter the global gene expression program to drive senescence.

## INTRODUCTION

Cellular senescence, a process characterized by prolonged and irreversible arrest of the cell cycle and associated with wide-ranging macromolecular damages and metabolic dysregulations (1, 2), is a hallmark of aging and a major contributor to diverse age-related diseases (ARDs) (3, 4). Among the various defining characteristics of cellular senescence is the extensive and multi-scale remodeling of the nuclear lamina (NL), a dense meshwork of lamin filaments and associated proteins at the nucleoplasmic face of the inner nuclear membrane that provides crucial structural support for the nucleus and helps connect the nuclear envelope to the chromatin inside (5). Senescence-related changes in NL structure, composition and the expression of its protein components have been known to significantly impact nuclear envelope integrity and nuclear morphology (6, 7). In particular, the expression of lamin B1 is widely known to be downregulated in many senescence-related contexts, leading to defects in nuclear shape and integrity as well as alternations in the expression of senescence-associated genes (8, 9). In addition, chromatin regions specifically associated with the NL, termed lamina-associated domains (LADs) (10), are also known to undergo large-scale genomic re-organization during senescence, altering NL–chromatin interactions that could potentially impact the global gene expression program to drive the aging process (11–14).

However, beyond these levels, it remains unclear if and how senescence-induced remodeling of NL composition (particularly the expression of lamins and various constituents of the NL-associated proteome) could quantitatively impact the spatio-temporal dynamics of the molecular interactions that occur at the NL, leading to alterations in the biophysical properties of the NL as well as of the overall nuclear envelope. Moreover, in light of the multi-scale ultrastructural changes known to occur to chromatin during cellular senescence (11, 15–17), the roles played by chromatin architecture in organizing and modulating such senescence-associated dynamics at the NL also remain to be investigated in a quantitative manner. To address these gaps, we choose as our model system the human retinal pigment epithelium (RPE) cells, a cell type crucial for proper vision, and the senescence of which is known to be a primary driver for the prevalent ocular ARD, age-related macular degeneration (AMD) (18, 19). However, despite its critical disease implications and wide adoption in aging studies, to date the spatio-temporal landscape of NL dynamics has not been systematically characterized in any RPE cellular system, in the context of senescence.

Herein, by combining live-cell single-molecule tracking/super-resolved mapping, imaging fluorescence correlation spectroscopy (imaging FCS) and transcriptomic profiling, we rigorously quantified the impact of cellular senescence on the NL dynamic landscape at the single-molecule level, using an inducible senescence model established in RPE cells that recapitulates key signatures of the senescence-associated phenotype and exhibits distinct alteration patterns in the expression of various NL components. In particular, we unveiled multi-modal changes in both the temporal dynamics as well as the spatial organization of NL mobility and binding interactions for two key mediators of NL–chromatin interactions, lamin B receptor (LBR) and emerin, in senescent RPE cells, hence revealing a previously overlooked facet of senescence-induced NL remodeling. Our findings also point to a potential mechanistic synergy between constitutive heterochromatin restructuring at the NL and dynamic attenuation of stable NL-binding interaction (particularly in the form of LBR binding “hotspots”) during senescence, leading to a loss of NL–heterochromatin contacts that could in turn have global repercussions for senescence-associated gene expression program.

## RESULTS

### Establishing a chemically inducible RPE cellular senescence model

We first sought to establish a method capable of robustly inducing senescence in RPE cells (specifically the ARPE-19 cell line) and recapitulating various features of the senescence-associated phenotype. To that end, we chose *tert*-butyl hydroperoxide (*t*BHP), a well-known oxidative stressor (20, 21) which has been demonstrated to induce senescence in other cell types such as human fibroblasts and dermal mesenchymal stem cells (22, 23). Upon treatment with daily doses of *t*BHP over a period of 4 days (see **Materials and Methods** for details), ARPE-19 cells showed drastically reduced cell proliferation as compared to untreated cells (**Fig. 1A**), in line with expectation from other cell types. Immunofluorescence staining and Western blotting both found that the levels of two key cell cycle arrest markers, p21 and phosphorylated retinoblastoma protein (phospho-Rb), exhibited a 2.7-fold increase and 94% decrease, respectively (**Fig. 1B** and **C**), indicating the activation of the p53–p21 and p16–phospho-Rb signaling pathways upon *t*BHP treatment to promote exit from the cell cycle into a less proliferative state. Moreover, senescence-associated β-galactosidase (β-gal) staining showed a drastic increase in the percentage of β-gal-positive ARPE-19 cells from an average of 10% to 73% upon *t*BHP treatment (**Fig. 1D**), suggesting that majority of the *t*BHP-treated cells have entered a senescent state. Similar reduction in cell proliferation, p21/phospho-Rb level alterations, and increase in β-gal-positive cells were observed upon treating ARPE-19 cells with doxorubicin (**Fig. S1A** to **C**), a drug known to also promote senescence via oxidative stress, in addition to other mechanisms (such as topoisomerase II inhibition-induced DNA damage (24)).

**Figure 1.**
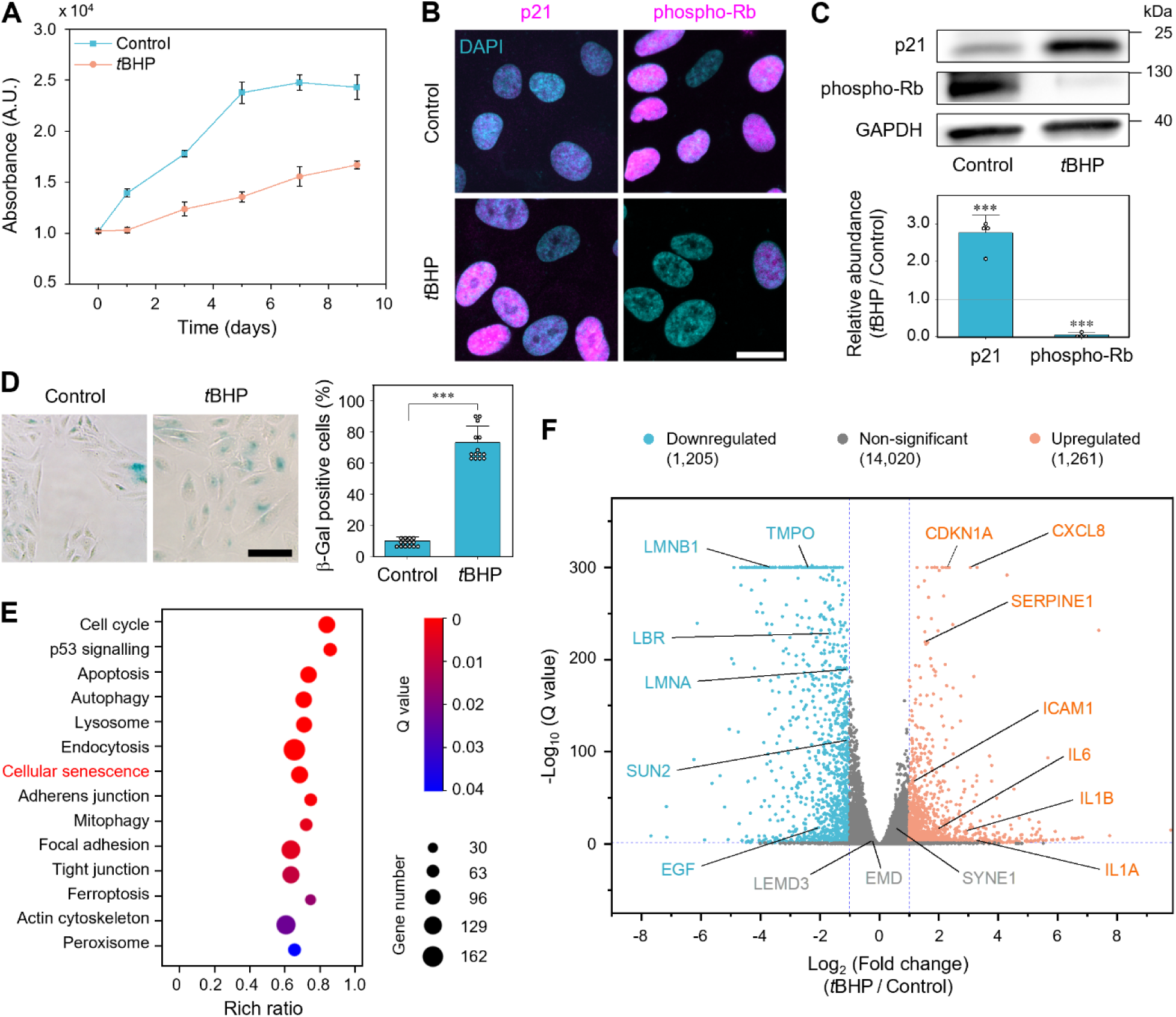
Modelling cellular senescence in RPE cells. **(A)** Cell proliferation profiles of *t*BHP-treated and untreated ARPE-19 cells (as measured by CCK-8 assay) over a period of 9 days; day 0 denotes the commencement of the first dose of *t*BHP treatment. **(B)** Representative confocal images of *t*BHP-treated and untreated ARPE-19 cells immunofluorescently stained for cell cycle arrest markers p21 or phospho-Rb; the cell nucleus is co-stained with DAPI. **(C)** Western blots for p21 or phospho-Rb in *t*BHP-treated and untreated ARPE-19 cells (top), as well as quantification of their relative abundances (normalized to GAPDH, bottom). **(D)** Representative bright-field images of *t*BHP-treated and untreated ARPE-19 cells stained for senescence-associated β-galactosidase (left), as well as the percentage of β-galactosidase-positive cells in each sample (right). **(E)** Kegg pathway enrichment analysis of RNA-seq data on *t*BHP-treated and untreated ARPE-19 cells, showing the top cellular processes-related hits upon *t*BHP treatment (including cellular senescence). **(F)** Differential expression analysis of genome-wide transcript levels in *t*BHP-treated and untreated ARPE-19 cells, showing significant up- or down-regulation of various SASP- and nuclear lamina-associated genes upon *t*BHP treatment. Scale bar: 20 µm (**B**); 100 µm (**D**). *n* = 3 (**A**), 4 (**C**) and 14 (**D**) independent replicates; data are represented as mean ± S.D..

Furthermore, whole-transcriptome RNA sequencing (RNA-seq) and KEGG pathway enrichment analysis (25) found senescence to be among the top cellular processes-related hits in ARPE-19 cells upon *t*BHP treatment (**Fig. 1E**). RNA-seq also identified significantly altered expression for a variety of senescent-associated secretory phenotype (SASP) genes, including those that encode interleukins (IL1A/B, IL6, CXCL8), epidermal growth factor (EGF), intercellular adhesion molecule 1 (ICAM1), and plasminogen activator inhibitor-1 (SERPINE1) (26, 27) (**Fig. 1F**). In addition, the upregulation of CDKN1A (which encodes p21) at the transcript level further corroborates our immunofluorescence and Western blotting results at the protein level (**Fig. 1B** and **C**). Collectively, these evidence demonstrate that *t*BHP treatment can successfully recapitulate a wide array of senescence-associated phenotypic features, thereby enabling us to model the senescent state in ARPE-19 cells in a robustly inducible manner. Similar enrichment of senescence-associated hits was again observed for doxorubicin-treated ARPE-19 cells (**Fig. S1D**); however, doxorubicin treatment exhibits substantially lower rich ratio (0.25 vs. 0.67 for *t*BHP treatment) and gene number (39 vs. 105 for *t*BHP treatment), as well as a much less significant Q value (0.029 vs. 0.00035 for *t*BHP treatment), suggesting that it mimics the senescent state in a less comprehensive and faithful manner. As such, we decided to use *t*BHP treatment for subsequent modeling and quantitative imaging/analyses.

### RPE cellular senescence remodels nuclear lamina composition and enlarges nuclear size

To assess the impact of *t*BHP-induced senescence on the NL in APRE-19 cells, we further performed GO cellular component analysis (28) on our RNA-seq data; indeed, nuclear lamina was found to be among the top nucleus-related hits associated with *t*BHP treatment (**Fig. 2A**). Moreover, a variety of NL-associated genes were significantly downregulated upon senescence induction in APRE-19 cells (**Fig. 1F**). Among them, lamin B1 (encoded by LMNB1 gene) is a major component that provides structural support for the NL and well-known to be downregulated in senescence (29), while lamina-associated polypeptide 2β (LAP2β, encoded by TMPO gene) and lamin B receptor (LBR) both bind lamins and facilitate the tethering of chromatin at the NL (30). On the other hand, several other key NL-associated genes such as lamin A/C (another major structural component of the NL, encoded by LMNA gene (31)) and SUN2 (an integral component of the LINC complex that mediates nucleo-cytoplasmic coupling (32)) exhibits only borderline downregulation, while the expression of emerin and MAN1 (members of the LEM-domain protein family also involved in providing structural anchorage by binding lamins and tethering chromatin at the NL, encoded by EMD and LEMD3 genes respectively (33)) as well as of nesprin-1 (another component of the LINC complex, encoded by SYNE1 gene (32)) remain largely unchanged upon *t*BHP treatment (**Fig. 1F**).

**Figure 2.**
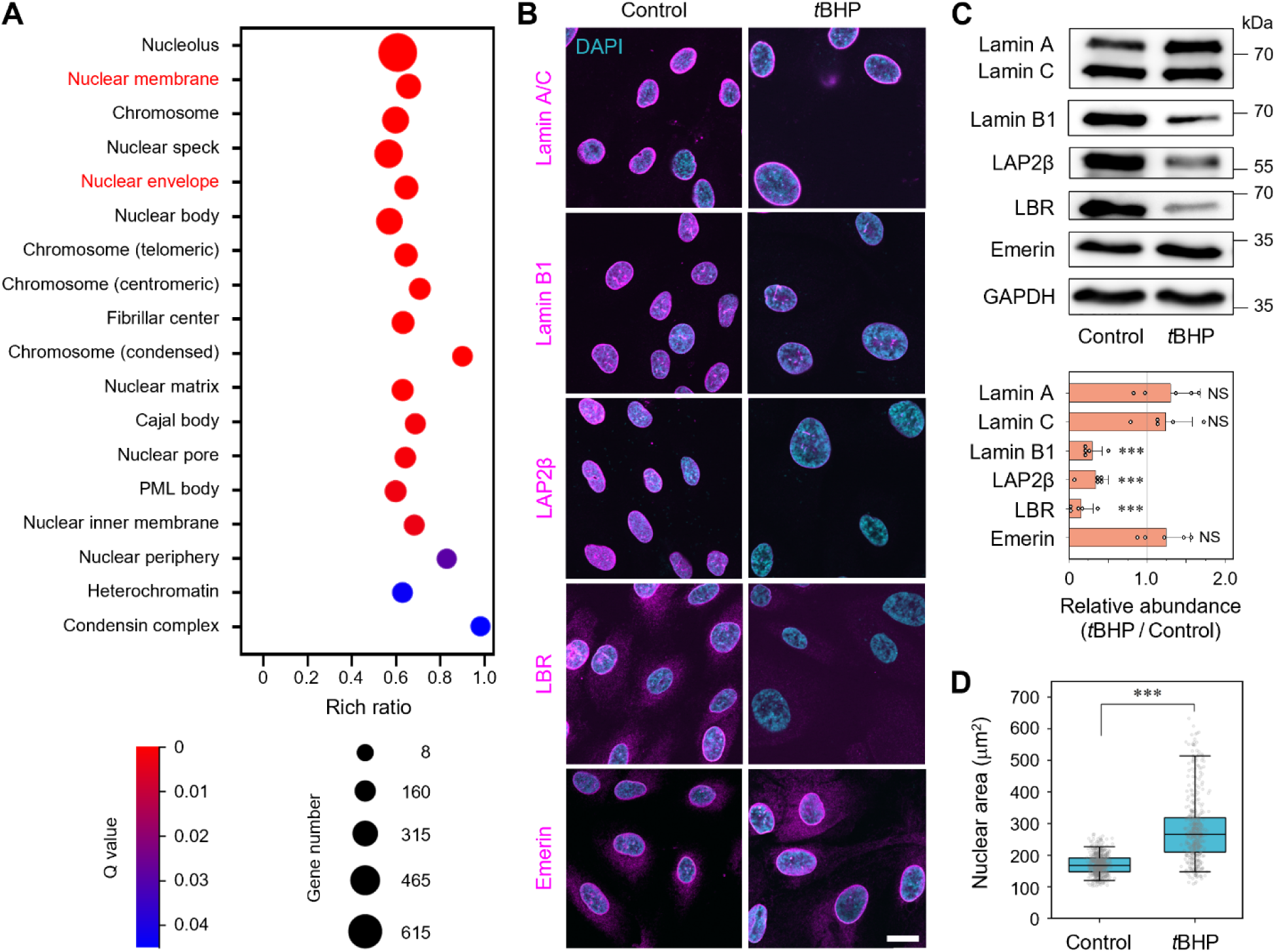
RPE cellular senescence is associated with distinct expression-level changes for key nuclear lamina components as well as nuclear size enlargement. **(A)** GO cellular component analysis of RNA-seq data on *t*BHP-treated and untreated ARPE-19 cells, showing the top nucleus-related hits upon *t*BHP treatment (including nuclear lamina, classified under both nuclear membrane and nuclear envelope categories). **(B)** Representative confocal images of *t*BHP-treated and untreated ARPE-19 cells immunofluorescently stained for each of 6 key components of the nuclear lamina: lamin A/C, lamin B1, LAP2β, LBR, and emerin; the cell nucleus is co-stained with DAPI. **(C)** Western blots for each of the 6 key nuclear lamina components in **(B)** in *t*BHP-treated and untreated ARPE-19 cells (top), as well as quantification of their relative abundances (normalized to GAPDH, bottom). **(D)** Nuclear areas of *t*BHP-treated and untreated ARPE-19 cells, quantified from DAPI-stained confocal images in **(B)**. Scale bar for **B**: 20 µm. *n* = 5 independent replicates for **C**, and 625 (control) and 449 (*t*BHP-treated) cells for **D**. Data are represented as mean ± S.D. for **C**, and median, 25^th^/75^th^ percentiles (box) and 5^th^/95^th^ percentiles (whiskers) for **D**.

To further confirm whether such transcript-level changes are transmitted to the protein level, we performed immunofluorescence staining as well as Western blotting for 6 key constituents of the NL-associated proteome in ARPE-19 cells (**Fig. 2B** and **C**). In line with the RNA-seq results, the levels of lamin B1, LAP2β and LBR were all significantly reduced by an average of 3.4, 2.9 or 6.4 folds respectively upon *t*BHP treatment, whereas the levels of lamin A/C and emerin remain largely unaffected. These changes, however, are not a result of non-senescence-associated cell cycle arrest, as evidenced by treating the cell with palbociclib (**Fig. S2**), a drug known to induce cell cycle arrest by hypo-phosphorylating Rb via the inhibition of cyclin-dependent kinase 4/6 (34). Finally, we also found that *t*BHP treatment leads to an average of 1.7-fold enlargement of nuclear size in APRE-19 cells (**Fig. 2D**), another well-known morphological feature associated with senescence (35, 36). This not only further validates our *t*BHP-induced senescence model, but also enables us to use nuclear size as a facile yardstick for selecting genuinely senescent cells for subsequent imaging analyses in the event of a non-homogeneously senescent cell population.

### LBR and emerin exhibit distinctly enhanced mobility indicative of weakened NL anchoring upon senescence induction

In light of the senescence-induced expression-level changes for various NL components in ARPE-19 cells, we next set out to interrogate how such expression-level changes could quantitatively impact their dynamic interactions with the NL as well as biophysical properties of the NL (and for that matter, of the nuclear envelope in general). To that end, we focused on LBR and emerin, both of which play key roles in bridging NL–chromatin interactions, but are known to bind preferentially to different lamins on the NL (LBR to lamin B1/2 vs. emerin to lamin A/C) as well as exhibit distinct patterns of expression-level alteration (significant reduction vs. minimal change) at both transcript and protein levels. We first quantified how their respective mobility on the NL is impacted by senescence using live-cell imaging FCS (37–39). In contrast to conventional single-point FCS that can only probe molecular mobility at one intracellular location at a time, imaging FCS (often performed under TIRF or HILO (40) illumination) can acquire multiple FCS measurements in a highly parallel fashion across an entire section of the NL; subsequent pixel-by-pixel autocorrelation analysis then generates a spatially resolved map of diffusion coefficient across the NL (**Fig. 3A**; see **Materials and Methods** for more details), hence making it ideally suited for quantifying global mobility changes upon senescence induction.

**Figure 3.**
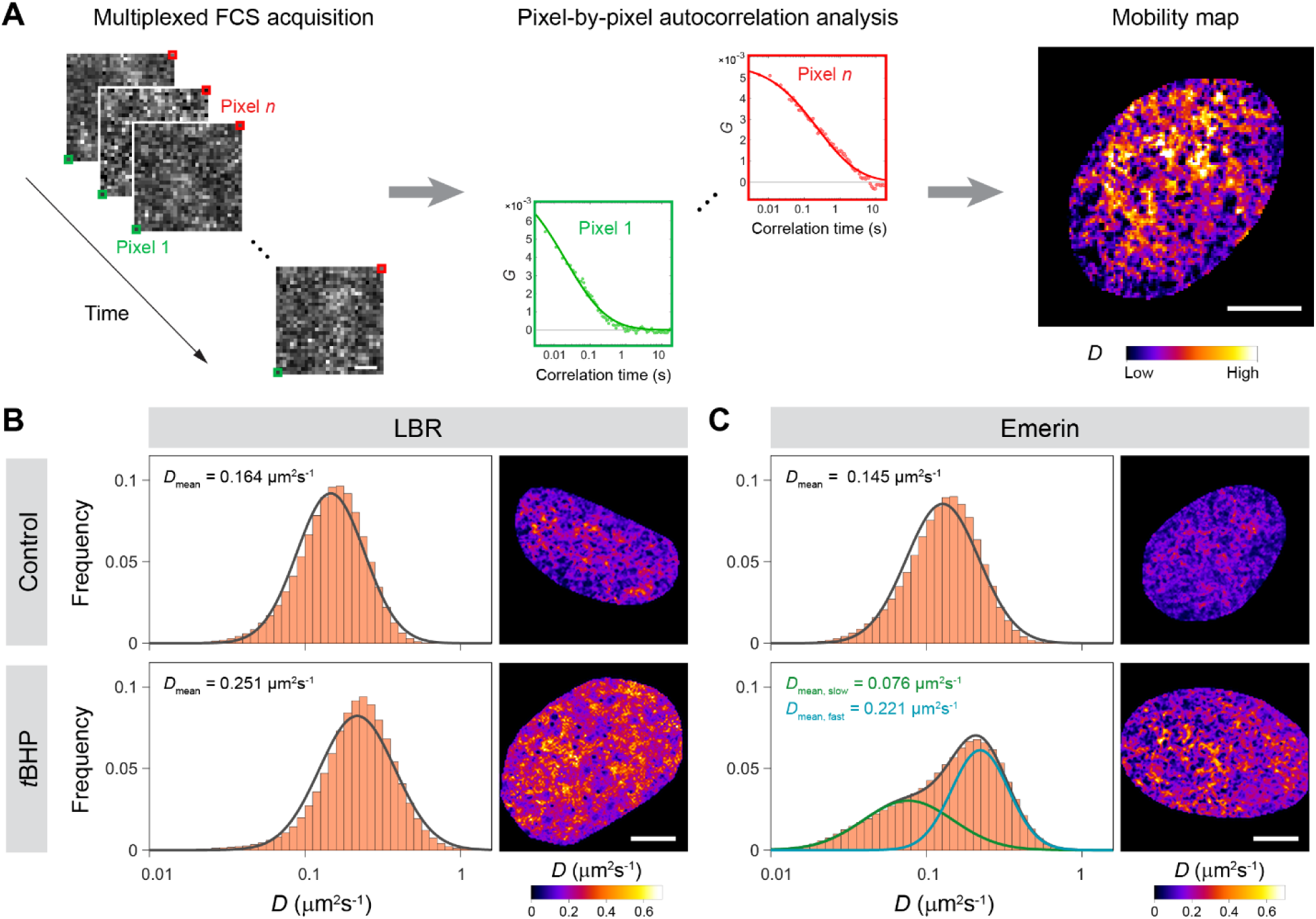
Live-cell imaging FCS reveals distinctly enhanced mobility for LBR and emerin on the nuclear envelope of senescent RPE cells. **(A)** Schematic of imaging FCS workflow for quantitative mapping of molecular mobility in live RPE cells. FCS measurements were acquired in a multiplexed fashion across the basal surface of the nuclear envelope under HILO illumination, followed by pixel-by-pixel autocorrelation analysis to generate a spatially resolved map of diffusion coefficient (*D*). **(B** and **C)** Histograms of diffusion coefficient (left column) as well as the corresponding mobility maps (right column) quantified from imaging FCS measurements on knock-in ARPE-19 cells endogenously expressing LBR-Halo **(B)** or Halo-emerin **(C)**, without or with *t*BHP treatment. Black curves in the histograms denote single-Gaussian fits, except for emerin in *t*BHP-treated cells where a double-Gaussian fit (with individual Gaussian components shown in green and cyan) is needed to satisfactorily fit the distribution. Scale bar: 5 µm. *n* = 63 (control) and 30 (*t*BHP-treated) cells for **B**, and 47 (control) and 31 (*t*BHP-treated) cells for **C**.

To do so, we generated two knock-in ARPE-19 cell lines expressing HaloTag-fused LBR or emerin at endogenous level using CRISPR/Cas9-mediated genome editing, and validated that they can be similarly induced into a senescent state with *t*BHP treatment while retaining the expression-level alteration patterns for the 6 key NL components as was observed for wild-type APRE-19 cells (**Fig. S3**). Performing imaging FCS under HILO illumination at a fast acquisition speed of 5.5 ms per frame on JF_549_-labeled LBR-Halo or Halo-emerin (**Movie S1**) at the basal surface of the nuclear envelope in these cells (**Fig. S4**), we found that both LBR and emerin exhibit significantly enhanced mobility upon *t*BHP treatment, as evidenced by a ∼53% increase in their diffusion coefficients (*D*, from mean values of 0.164 µm^2^s^-1^ and 0.145 µm^2^s^-1^ to 0.251 µm^2^s^-1^ and 0.221 µm^2^s^-1^, respectively) (**Fig. 3B** and **C**), indicating that senescence weakens NL’s structural anchoring on the inner nuclear membrane (on which LBR and emerin undergo lateral diffusion) and makes it more fluid. Intriguingly, in contrast to the uni-modal distribution of diffusion coefficient observed for LBR, the diffusion coefficient of emerin in senescent ARPE-19 cells exhibits a bi-modal distribution, consisting of an additional minor fraction with substantially reduced mobility (with a mean *D* of 0.076 µm^2^s^-1^) on top of the aforementioned faster-diffusing fraction. Moreover, the mobility maps of both LBR and emerin exhibit significant spatial heterogeneities, which become more pronounced in senescent cells. In particular, the numerous low-mobility regions across the NL potentially correspond to sites where LBR or emerin has a higher propensity to interact with both the NL and the chromatin in the vicinity for longer durations, likely with functionally relevant consequences. Finally, to ascertain that these changes are indeed a consequence of senescence-associated NL remodeling, we again performed similar imaging FCS measurements on knock-in ARPE-19 cells treated with either palbociclib or two additional drugs known to alter chromatin accessibility, TSA (a pan-histone deacetylase inhibitor that decondenses chromatin globally (41)) and QC6352 (a specific inhibitor of KDM4 histone demethylase that induces localized chromatin condensation (42–44)). All of these treatments were found to perturb the mobilities of both LBR and emerin only marginally as compared to *t*BHP, both in terms of the distributions of their diffusion coefficients and their spatial heterogeneities across the NL (**Fig. S5**), hence verifying that the distinct mobility enhancements observed for LBR and emerin are not a consequence of non-senescence-associated cell cycle arrest or chromatin accessibility changes in RPE cells.

### Single-molecule tracking and STAR mapping unveil senescence-induced spatio-temporal re-organization of NL-binding dynamics for LBR and emerin

In light of the highly heterogeneous mobilities observed for LBR and emerin, we next investigated how senescence impacts the organization and dynamics of their binding interactions at the NL in both space and time. To do so, we resorted to live-cell single-molecule tracking (SMT), which is known to be more suitable than FCS for probing dynamic processes that occur on longer timescales than diffusion, especially binding (45). In contrast to the much faster acquisition speed used for imaging FCS, we performed SMT on JF_549_-labeled LBR-Halo or Halo-emerin in our knock-in APRE-19 cells at 300 ms per frame, in order to blur out most of the rapidly diffusing molecules and focus instead on their binding events on the NL (**Movie S2**). To further eliminate potentially contaminating diffusion events from the NL-binding trajectories (**Fig. 4A**), we implemented a previously devised strategy capable of unambiguously discriminating binding from diffusion (46), and subjected the resulting NL-binding trajectories to survival probability analysis (47). The survival time distributions we obtained are better fitted with a three-state model as compared to a two-state model (**Fig. 4B**), indicating the existence of three temporally distinct NL-binding modes for both LBR and emerin: a transient binding mode (with a mean residence time (τ) of 0.67–0.73 s, during which they likely sample the NL nonspecifically), and two stable binding modes (with mean residence times of 3.0–3.1 s for the shorter-lived mode and 14.6–14.7 s for the longer-lived mode, during which specific interactions with NL targets likely occur) (**Fig. 4C**). As a validation, similar SMT measurement on NRF2, a transcription factor involved in oxidative stress response but does not bind the NL under normal conditions (48), exhibits only 2 binding modes with residence times distinctly different from those of LBR and emerin (**Fig. S6**), hence ruling out the possibility of the binding modes we observed to be measurement- or analysis-related artefacts.

**Figure 4.**
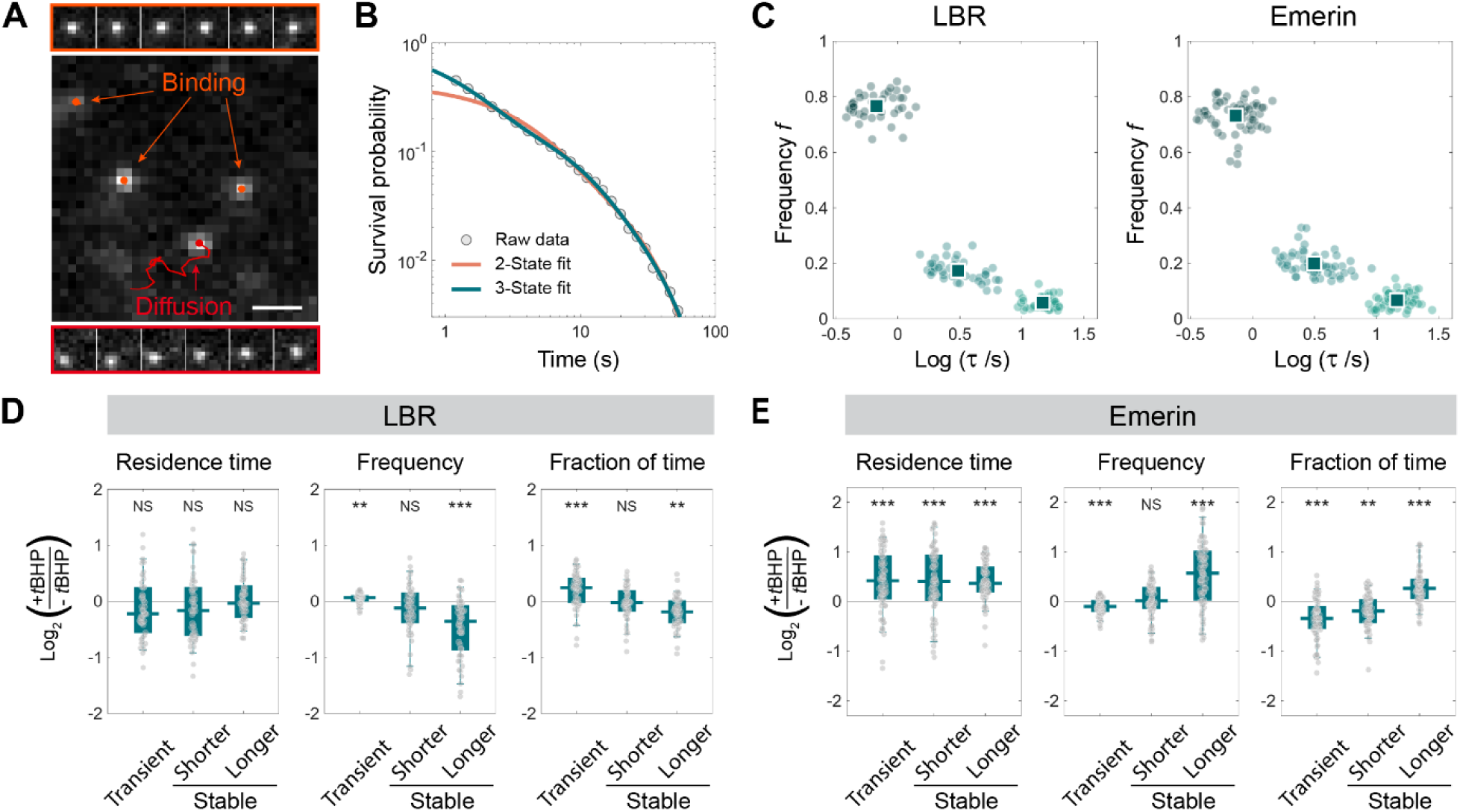
LBR and emerin exhibit distinct, multi-modal alterations in NL-binding dynamics induced by RPE cellular senescence. **(A)** A representative frame from SMT movies of LBR-Halo or Halo-emerin in live ARPE-19 cells, showing both diffusion and binding events on the NL (with an example of each shown above and below, respectively); individual trajectories of bound (orange) or diffusing (red) molecules are superimposed. Only binding events are subjected to subsequent analyses. **(B)** A typical survival probability distribution derived from SMT binding trajectories in **(A)**, fitted with either a two-state model (orange) or a three-state model (green). **(C)** Scatter plots of residence time (τ) and binding frequency (*f*) associated with each mode of NL-binding resolved for LBR (left) or emerin (right) from SMT measurements. **(D** and **E)** Box-and-whisker plots of fold changes (on log_2_ scale) in residence time, binding frequency and fraction of time bound associated with each mode of NL-binding for LBR **(D)** or emerin **(E)** upon *t*BHP treatment. Scale bar for **A**: 1 µm. In **C**–**E**, each dot denotes a single cell, with *n* = 43 (LBR) and 57 (emerin) cells for **C**, 43 (control) and 49 (*t*BHP-treated) cells for **D**, and 57 (control) and 67 (*t*BHP-treated) cells for **E**. Squares in **C** denote mean values, whereas in **D** and **E** data are represented as median, 25^th^/75^th^ percentiles (box) and 5^th^/95^th^ percentiles (whiskers).

Upon inducing senescence with *t*BHP, LBR and emerin exhibit distinct alteration patterns in their NL-binding dynamics, in terms of residence time, binding frequency (*f*), as well as the fraction of time bound under each mode (*F*, defined as the normalized product of τ and *f* for each binding mode (46)) (**Fig. 4D** and **E**). Specifically, senescence leads to significant increases in the residence times of all three binding modes (by an average of 29–34%) for emerin but not for LBR, echoing the substantially less mobile fraction observed uniquely for emerin from imaging FCS measurements (**Fig. 3C**). LBR in senescent ARPE-19 cells also exhibited a marked shift in its molecular partitioning from stable binding (particularly the longer-lived mode) towards transient binding, as manifested by both binding frequency and fraction of time bound; on the other hand, emerin exhibited the opposite trend where longer-lived stable binding becomes more favored at the expense of transient binding, upon senescence induction. In contrast, these alteration patterns were either not observed or occurred to a much less extent in cells treated with either palbociclib or TSA (**Fig. S7**), again demonstrating that they are not a consequence of non-senescence-associated cell cycle arrest or global changes in chromatin accessibility. Intriguingly, QC6352 treatment shifted the molecular partitioning of LBR towards longer-lived stable binding (*i.e.* opposite of the trend observed for *t*BHP treatment) (**Fig. S7A**), hence suggesting a potential role played by localized chromatin decondensation in modulating the NL-binding dynamics of LBR.

Moreover, given that the mobility maps obtained earlier from imaging FCS are intrinsically diffraction-limited, we further leveraged a recently developed strategy, termed STep Accumulation Reconstruction (STAR), capable of super-resolved density mapping of the spatial distribution of intranuclear binding (46). Conceptually akin to how single-molecule localization microscopy techniques make use of the positions of individual molecules to reconstruct a super-resolution image, STAR uses the positions of individual steps in SMT trajectories to inform on the frequency with which binding events occur at each intranuclear location. Applying STAR mapping to the SMT trajectories of LBR and emerin from live APRE-19 cells in a binding-mode-specific manner (**Fig. 5A** and **B**), we found that their NL-binding events (for both transient and stable modes) exhibited a spatially heterogeneous distribution resembling the mobility maps from imaging FCS measurements, and preferentially occur at numerous distinct loci scattered across the NL. In particular, stable binding tends to be clustered into “hotspots” at specific loci where multiple binding events occur sequentially and repeatedly, as opposed to transient binding which occurs only as unclustered, singular binding events.

**Figure 5.**
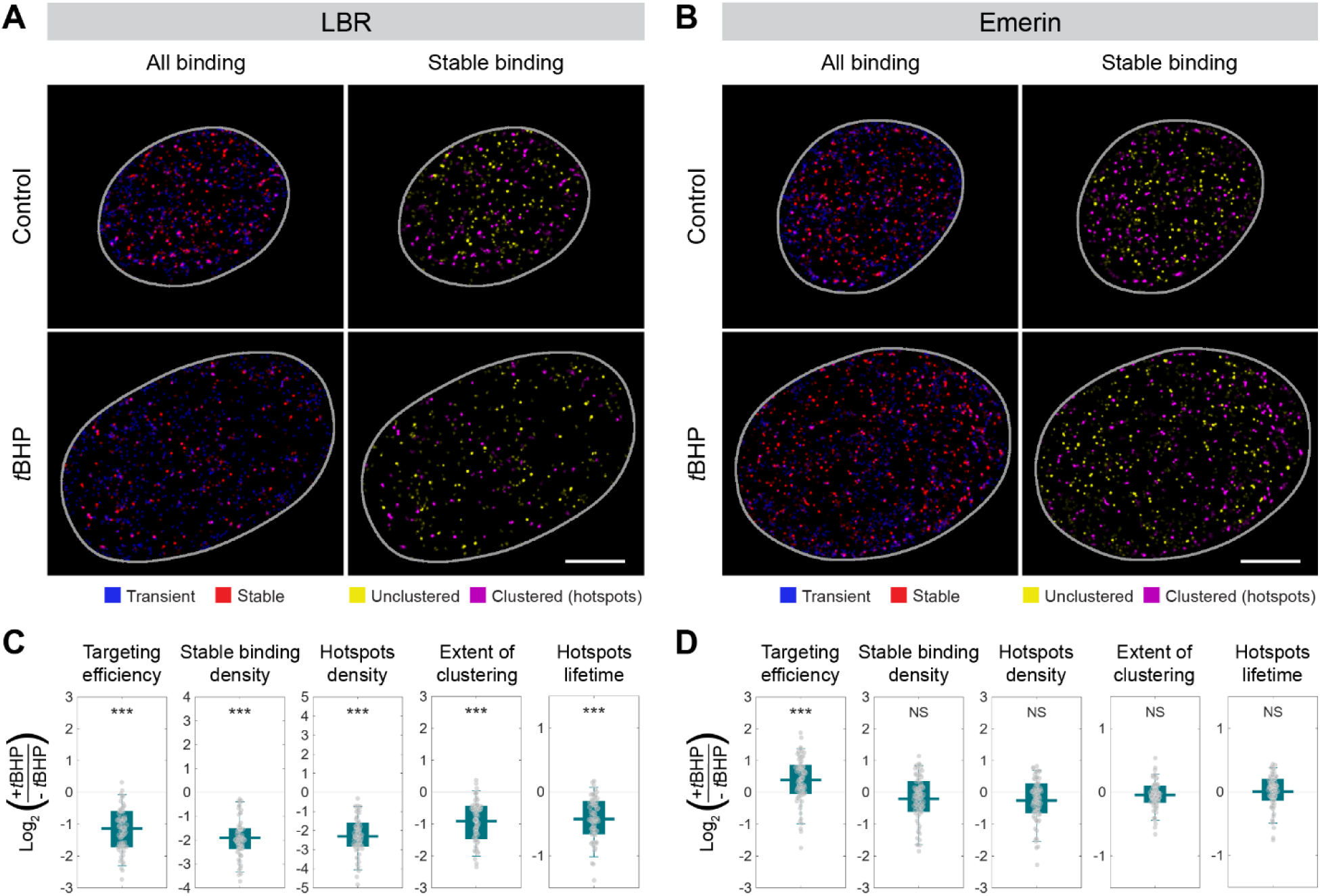
STAR mapping unveils differentially re-organized NL-binding landscape for LBR and emerin in both space and time. **(A** and **B)** Super-resolved NL-binding maps for LBR **(A)** or emerin **(B)** in live ARPE-19 cells without or with *t*BHP treatment, showing either all binding modes combined (left column) or stable binding only (right column, categorized into unclustered binding events or binding clusters (*i.e.* “hotspots”)). The nuclear boundary is delineated in grey. **(C** and **D)** Box-and-whisker plots of fold changes (on log_2_ scale) in targeting efficiency, stable binding events density, stable binding hotspots density, extent of clustering for stable binding, as well as binding hotspots lifetime for LBR **(C)** or emerin **(D)** upon *t*BHP treatment. Scale bar for **A**–**B**: 5 µm. In **C**–**D**, each dot denotes a single cell, with *n* = 45 (control) and 60 (*t*BHP-treated) cells for **C**, and 61 (control) and 69 (*t*BHP-treated) cells for **D**; data are represented as median, 25^th^/75^th^ percentiles (box) and 5^th^/95^th^ percentiles (whiskers).

Importantly, upon senescence induction with *t*BHP, the spatial distributions of NL-binding for LBR and emerin were found to undergo distinct and multi-modal re-organization (**Fig. 5C** and **D**). To begin with, the relative proportion of stable binding loci was reduced by an average of 55% for LBR, a trend quantifiable using the metric targeting efficiency (defined as the ratio between the NL space explored by stable binding versus that by diffusion and transient binding combined). Moreover, the density of stable binding events (per unit area on the NL) for LBR was also significantly reduced by an average of 74% upon *t*BHP treatment. Both of these findings are in line with the senescence-induced downregulation observed for LBR as well as its NL-binding partner, lamin B1 (**Fig. 2B** and **C**). In addition to reductions in sheer number and targeting efficiency, the spatial distribution of stable binding events was also re-organized for LBR, as evidenced by the density of stable binding hotspots as well as their extent of clustering (defined as the ratio between stable binding events that cluster into hotspots vs. those that exist in unclustered form), both of which exhibited marked reductions by 80% and 43%, respectively. These indicate that the downregulation of LBR in senescent cells not only makes it more difficult to target specific NL sites for stable binding, but also makes stable NL-binding events spatially more sparse and hence less likely to congregate into hotspots where they can occur sequentially and repeatedly, likely for functionally relevant purposes. In contrast, the densities of stable binding events and hotspots as well as the extent of clustering for emerin all remain largely unchanged in senescent cells, again consistent with its unaltered expression level as well as that of its NL-binding partner, lamin A/C (**Fig. 2B** and **C**). Emerin’s targeting efficiency, however, increases substantially by an average of 31%, likely a consequence of the shift in its molecular partitioning from transient binding towards stable binding (**Fig. 4E**). Finally, in the temporal domain, the lifetime of stable binding hotspots (defined as the total duration spanned by all binding events within each hotspot) for LBR is reduced by an average of 26% (in line with fewer stable binding events being clustered into hotspots), while that for emerin remains unaffected despite its longer residence times for all binding modes. This indicates that the residence times of binding account for a much smaller fraction of the total duration of each binding hotspot as compared to the gap times between successive binding events that temporally adjoin them into a hotspot, further pointing to the potential functional significance of such clustering behavior in both space and time.

### Correlative mapping links senescence-induced re-organization of LBR dynamics with constitutive heterochromatin restructuring

In order to further understand the mechanistic underpinning of the dynamic changes we have observed, we noted that QC6352 treatment led to an effect on LBR’s NL-binding dynamics opposite to that of *t*BHP-induced senescence (**Fig. S7A**). In fact, the KDM4 family of demethylases inhibited by QC6352 removes histone mark H3K9me3 (42), which is specifically recognized and bound by the HP1α protein instrumental in condensing DNA into constitutive heterochromatin (49), which is known to be largely associated with the NL (10). Hence, we hypothesize that the reverse process of decondensing HP1α-rich constitutive heterochromatin could play a key role in re-organizing the NL-binding dynamics of LBR, which is itself known to interact directly with heterochromatin and tether it to the inner nuclear membrane (30). Indeed, restructuring of chromatin organization is known to occur during senescence across different scales, ranging from changes in epigenetic modifications at the nucleosome level, to altered interaction landscape within topologically associating domains and switching of A/B compartments at the megabase scale, and all the way to macroscale detachment of LADs from the NL (11, 50, 51).

To test this hypothesis, we performed correlative mapping by sequentially integrating imaging FCS with confocal microscopy to correlate, within the same ARPE-19 cell, the mobility map of LBR with the spatial organization of constitutive heterochromatin at the NL. In addition to the prevalently existing small foci of constitutive heterochromatin (marked by HP1α), nonsenescent ARPE-19 cells contain a small number of much larger foci corresponding to highly condensed heterochromatin domains at the NL, which become dispersed in *t*BHP-treated senescent cells where only smaller foci exist (**Figs. 6A** and **S8**). Upon superposition with the LBR mobility map measured at the same NL basal surface, we segmented the HP1α foci and performed radial profile analysis of HP1α intensity and LBR mobility centered on each focus (see **Materials and Methods** for details) (**Fig. 6B** and **C**). We found that the larger HP1α foci (with area >0.4 μm^2^) present only in nonsenescent cells are predominantly anti-correlated with LBR mobility on the NL. This trend, however, is not discernible in nonsenescent cells if all HP1α foci are pooled together regardless of their size, or in *t*BHP-treated senescent cells that contain only smaller HP1α foci. In both of these cases, the fractions of correlated, non-correlated, and anti-correlated HP1α foci observed closely match those obtained from applying the same analysis to simulated distributions of HP1α and LBR mobility, generated independently using relevant parameters derived from experimental imaging data (**Figs. 6C and S9**). This similarity indicates that the spatial relationship between HP1α foci and LBR mobility in both cases is consistent with a distribution expected of the absence of spatial coupling. Collectively, these results indicate that highly condensed domains of HP1α-rich constitutive heterochromatin are strongly correlated with regions of reduced LBR mobility on the NL, which most likely correspond to sites where LBR undergoes temporally extended interactions with the NL and heterochromatin via stable binding. The dispersal of these large heterochromatin domains during senescence diminishes these extended interactions, leading to a loss of NL– heterochromatin contacts that could potentially have functionally important consequences (*e.g.* alterations to global gene expression program known to be associated with senescence).

**Figure 6.**
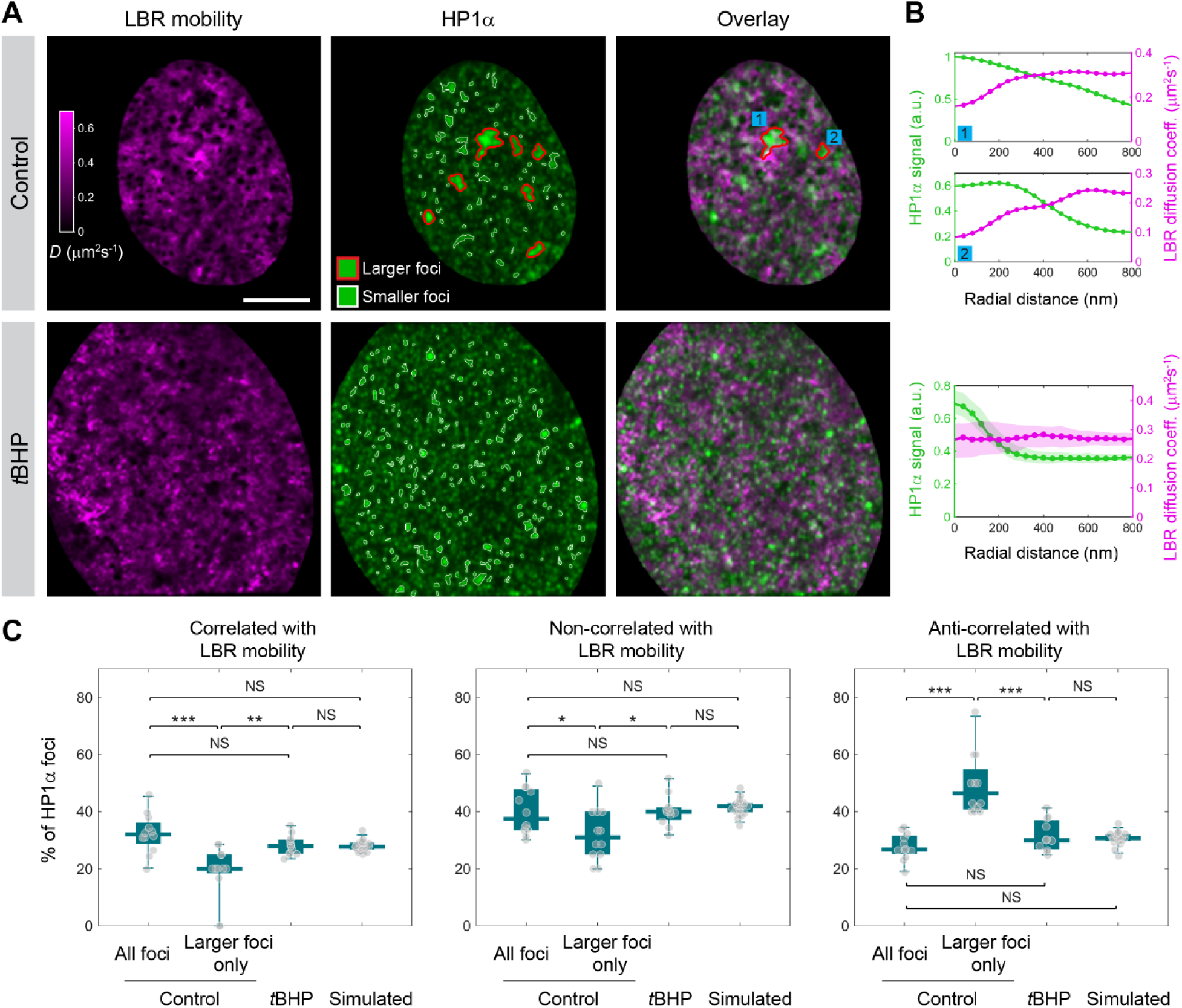
Senescence-induced re-organization of LBR dynamics on the nuclear lamina correlates with constitutive heterochromatin restructuring. **(A)** Representative correlative maps from knock-in ARPE-19 cells without (top row) or with (bottom row) *t*BHP treatment, showing the mobility map of LBR-Halo (labeled with JF_549_ and measured with imaging FCS) and the distribution of constitutive heterochromatin foci (as marked by immunofluorescently labeled HP1α and imaged with confocal microscopy) across the same NL basal surface in the same cell, as well as an overlay of the two. Larger HP1α foci (with area >0.4 μm^2^) identified through image segmentation are delineated in red, while smaller foci are delineated in white. **(B)** Correlative radial profile analysis between HP1α signal and LBR mobility on the two cells shown in **(A)**, either for the two larger HP1α foci indicated (for control cell, top) or averaged across all smaller foci (for *t*BHP-treated cell, bottom). **(C)** Box-and-whisker plots of the percentage of HP1α foci at the NL that are either correlated (left), non-correlated (middle) or anti-correlated (right) with LBR mobility in APRE-19 cells either without or with *t*BHP treatment, as well as in simulated distributions generated using relevant parameters derived from experimental imaging data. Only the larger HP1α foci in nonsenescent cells are predominantly anti-correlated with LBR mobility. Scale bar for **A**: 5 µm. In **C**, each dot denotes a single cell, with *n* = 12 (control), 11 (*t*BHP-treated) and 20 (simulated) cells; data are represented as median, 25^th^/75^th^ percentiles (box) and 5^th^/95^th^ percentiles (whiskers).

## DISCUSSION

In this study, we leveraged a multi-disciplinary approach and performed, to our knowledge, the first single-molecule characterization of the spatio-temporal dynamic landscape of the NL in a senescence-related human cell system. In addition to robustly recapitulating key signatures of the senescence-associated phenotype, our RPE cellular model revealed distinct alteration patterns for the expression of various key components of the NL, corroborated across transcript and protein levels. More importantly, our live-cell SMT and STAR mapping on LBR and emerin quantified a variety of spatio-temporal parameters of their dynamic interactions with the NL and chromatin, many of which were measured for the first time (*e.g.* density of binding events/hotspots, targeting efficiency, extent of clustering, and hotspots lifetime). Complementary measurements by imaging FCS further revealed their globally enhanced mobility, indicating that senescence weakens NL’s structural anchoring on the inner nuclear membrane and makes it more fluid. Moreover, the spatial distributions of the mobility and NL-binding for LBR and emerin, as unveiled by these measurements, are both highly heterogeneous across the NL, with numerous low-mobility regions or binding hotspots at which prolonged and repeated interactions with the NL and chromatin preferentially occur. Much of these quantitative information is hitherto unknown in any senescence-related context, thereby shedding light onto a previously inaccessible dimension of senescence-induced remodeling and re-organization of the NL at the single-molecule level.

Notably, the differential alterations in multiple spatio-temporal parameters of NL-binding dynamics observed for LBR and emerin (*e.g.* densities of stable binding events and hotspots, and hotspots lifetime) are in line with the distinct senescence-associated changes in both their own expression as well as the expression of their respective lamin binding partners on the NL (lamin B1 vs. lamin A/C). Moreover, LBR’s marked reduction in the extent of clustering for stable binding events indicates that its downregulation in senescent cells not only makes it more difficult to target specific NL sites for stable binding, but also makes stable binding events spatially more sparse and hence less likely to congregate into hotspots where more extended interactions can occur. Such spatial re-organization, as a strategy for senescence-induced remodeling of NL dynamic landscape, is complemented in the temporal domain by the significant increases in the binding residence times of emerin, which echo with its low-mobility fraction observed in imaging FCS measurements. These differential alterations in turn shed light on the distinct dynamic mechanisms potentially employed by LBR and emerin in modulating NL–chromatin interactions, despite their shared 3-mode binding and similar binding residence times (**Fig. 4C**).

In light of the fact that LBR directly interacts with heterochromatin and tethers it to the NL (as opposed to emerin which interacts with heterochromatin indirectly through bridging proteins such as barrier-to-autointegration factor (BANF1) (52), our correlative mapping and analysis further showed that regions of reduced LBR mobility on the NL are strongly correlated with highly condensed domains of constitutive heterochromatin at the NL, which become dispersed into much smaller foci upon senescence induction. Such a large-scale architectural re-organization weakens NL’s anchoring effect on heterochromatin, and leads to a loss of NL–heterochromatin contacts mediated by LBR (likely through stable binding interactions). These could in turn have functionally important consequences, especially in light of the recent findings that the loss of NL-heterochromatin tethering capabilities conferred by LBR (as well as by LAP2β) not only induces global detachment of heterochromatin from the NL, but also causes alterations in 3D chromatin organization, loss of epigenetic marks as well as dysregulation of gene expression profiles (30). On the other hand, artificially anchoring heterochromatin to the NL has been shown to mitigate senescence-associated phenotype and silence senescence-associated gene expression (53), further underscoring their relevance to our cellular senescence model.

Collectively, our findings point to a mechanistic model for senescence-induced re-organization of the NL dynamic landscape in both space and time, mediated by LBR and coupled with constitutive heterochromatin restructuring (**Fig. 7**). LBR is known to use a C-terminal transmembrane domain to span the inner nuclear membrane on which it undergoes lateral diffusion, while a long nucleoplasmic N-terminal region (containing a globular domain, a Tudor domain and a flexible hinge) mediates its various intranuclear interactions, chief among which being NL– heterochromatin contact (via HP1α) (49, 54, 55). During cellular senescence in RPE cells, large condensed domains of constitutive heterochromatin tethered to the NL get dispersed into smaller foci with weaker anchoring effect, while the downregulation of LBR as well as of lamin B1 on the NL significantly reduces both the temporal frequency and spatial density of stable binding of LBR (especially in the form of “hotspots” where multiple stable binding events spatio-temporally cluster to effect more extended and prolonged interactions with the NL and heterochromatin). Emerin, on the other hand, does not undergo such expression-level change or diminished stable binding interactions with the NL. The synergistic actions of such architectural re-organization, physical de-anchoring and dynamic attenuation lead to a less anchored inner nuclear membrane as well as a loss of LBR-mediated NL–heterochromatin contacts, which could in turn cause senescence-associated gene loci originally repressed within NL-tethered heterochromatin to detach from the NL and drive their upregulation during RPE cellular senescence. Indeed, comparison with an existing database of LADs (profiled in RPE-1 cells with pA-DamID (56)) showed that 25 (or 50%) of the 50 most upregulated genes detected in our *t*BHP-induced senescence model fall within the genomic boundaries of known LADs (**Fig. S10**), much higher than the average fraction of 13–18% of all genes typically found in a mammalian genome (57). As such, our model integrates the spatio-temporal alterations in NL dynamics uncovered in this study with the functional domain organization of LBR as well as the large-scale restructuring of chromatin architecture known to occur during cellular senescence, thereby shedding uniquely valuable insights into the genomic consequences of their mechanistic interplay.

**Figure 7.**
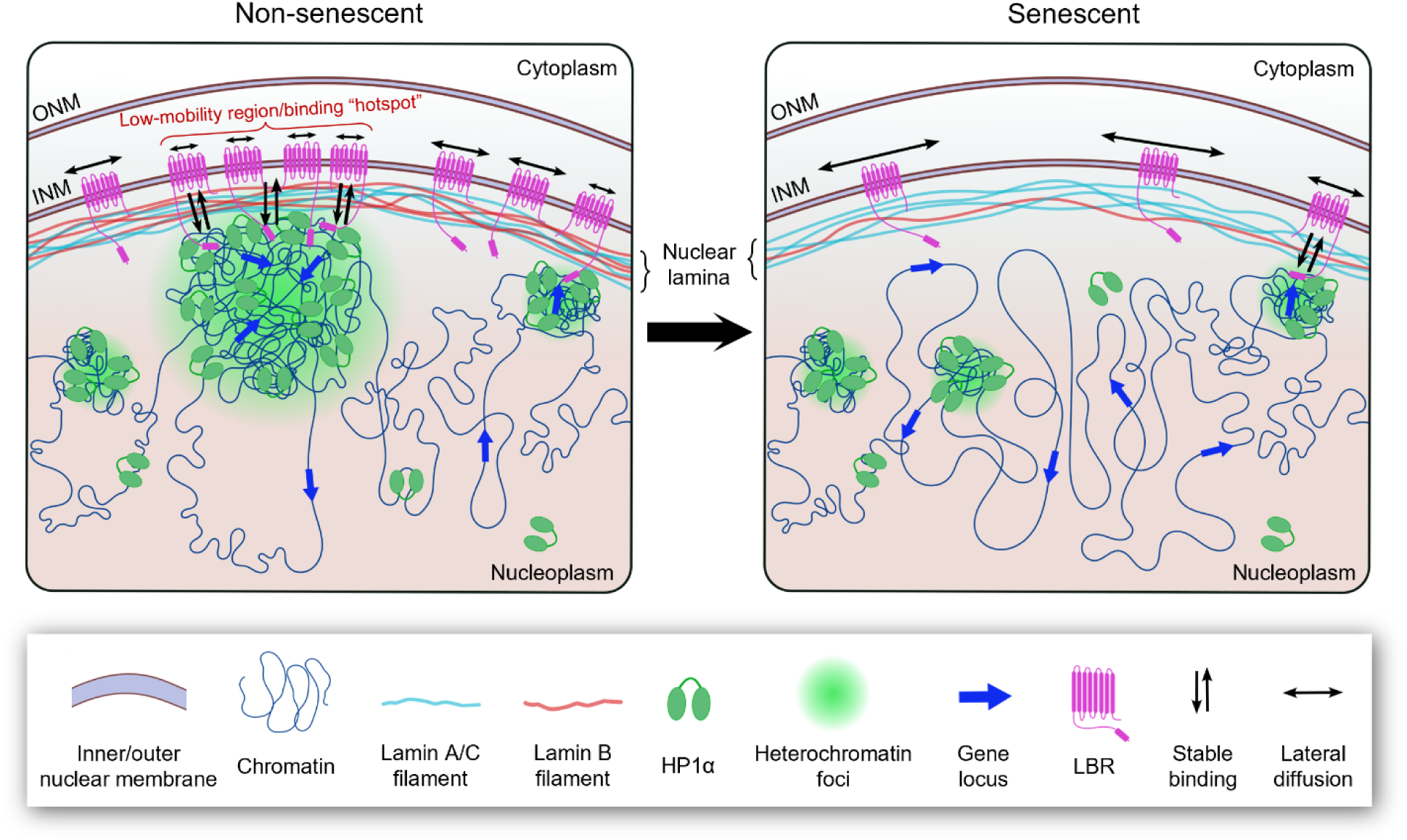
A mechanistic model for senescence-induced spatio-temporal re-organization of the nuclear lamina dynamic landscape in RPE cells. During cellular senescence, the dispersal of large condensed domains of HP1α-rich constitutive heterochromatin (originally tethered to the NL via LBR’s long N-terminal region) is accompanied by the downregulation of LBR and lamin B1, which significantly reduces LBR’s stable binding interactions on the NL in both spatial density as well as temporal frequency (particularly in the form of “hotspots” where multiple stable binding events cluster). Such architectural re-organization, physical de-anchoring and dynamic attenuation synergistically lead to a loss of LBR-mediated NL–heterochromatin contacts, which could in turn cause senescence-associated genes originally repressed within NL-tethered heterochromatin to detach from the NL and potentially drive their upregulation. The less anchored inner nuclear membrane also allows LBR to undergo faster diffusion, enhancing its mobility and further reducing its propensity for mediating prolonged NL–heterochromatin interactions.

Overall, the findings from our study demonstrate the re-organization of NL dynamic landscape at single-molecule level as a critical yet previously under-explored facet of senescence-induced NL remodeling in human RPE cells, which could be generally applicable to other cell types. The distinctive spatio-temporal correlation between LBR mobility/binding on the NL and constitutive heterochromatin restructuring unveiled in this study also provides first-of-its-kind quantitative evidence for the role of higher-order chromatin architecture in regulating NL dynamics, paving the way for further explorations into its diverse biological consequences. For example, the precise functional implications of emerin’s prolonged binding residence times in senescent cells (in line with its low-mobility fraction observed in imaging FCS measurements) warrant a more in-depth investigation, especially in the context of its role in providing structural anchorage at the NL through indirectly mediating NL–heterochromatin interactions. Comprehensive LADs mapping in our RPE cellular model (using *e.g.* ChIP-seq, CUT&Tag or DamID) as well as systematic colocalization (using *e.g.* DNA FISH) between the intranuclear loci of various genes upregulated during senescence and the low-mobility regions/binding hotspots on the NL will further allow us to correlate the altered NL dynamic landscape in physical space with their genomic targets in sequence space. Finally, our RPE cellular model as well as the quantitative approaches developed in this study could also be applied to probe other key components of the NL-associated proteome, as well as to diseased samples (*e.g.* AMD) in which RPE cellular senescence is known to be implicated, so as to understand how such aberrations in NL dynamic landscape could underpin various senescence-induced pathologies *in vivo*.

## MATERIALS AND METHODS

### Cell culture

ARPE-19 cells were grown in HyClone™ Dulbecco’s Modified Eagle Medium (DMEM)/F12 1:1 mixture (SH30023.01, Cytiva) supplemented with 10% (v/v) fetal bovine serum (FBS, 10500064, Life Technologies), 100 units/ml penicillin and 100 µg/ml streptomycin (Pen-Strep, 15140122, Life Technologies), and cultured at 37 °C under 5% CO_2_ in a humidified incubator. Prior to imaging, cells were seeded in a 35-mm dish with a 10-mm glass-bottom microwell and no. 1.5 coverglass (P35G-1.5-10-C, MatTek).

### Senescence induction and drug treatments

For *tert*-butyl hydroperoxide (*t*BHP) treatment, ARPE-19 cells were treated with 100 µM *t*BHP (458139, Sigma-Aldrich) in serum-free DMEM/F12 medium for 2 hours, and then washed with phosphate-buffered saline (PBS, 70011044, Gibco), and incubated with supplemented DMEM/F12 medium for at least 2 hours for the cells to recover. Cells were treated in the same fashion twice a day for four consecutive days, after which they were allowed to recover in culture for an additional 72 hours before subsequent experiments. For doxorubicin treatment, cells were exposed to 50 nM of doxorubicin hydrochloride (S1208, Selleckchem) in supplemented DMEM/F12 medium for 48 hours, after which the drug was removed, and the cells were cultured in supplemented DMEM/F12 medium for another 72 hours before subsequent experiments. For palbociclib treatment, cells were treated with 1 µM palbociclib (S1116, Selleckchem) in supplemented DMEM/F12 medium for 48 hours. For TSA treatment, cells were treated with 250 nM TSA (T1952, Sigma-Aldrich) in supplemented DMEM/F12 medium for 24 hours. For QC6352 treatment, cells were treated with 1 µM QC6352 (HY-104048, MedChemExpress) in supplemented DMEM/F12 medium for 72 hours.

### Cell proliferation assay

Cell Counting Kit-8 (CCK-8) (C0005, TargetMol) was applied to ARPE-19 cells in culture according to manufacturer’s protocol and incubated for 4 hours at 37 °C under 5% CO_2_ in a humidified incubator, after which the absorbance (at 490 nm) of the culture medium was measured with a 96-well microplate reader (Sunrise, Tecan).

### β-galactosidase assay

ARPE-19 cells were cultured in 6-well plates and stained with Senescence β-Galactosidase Staining Kit (9860, Cell Signaling Technology) according to manufacturer’s protocol overnight at 37 °C in a dry incubator, and then imaged with an EVOS M5000 brightfield light microscope (Thermo Fisher Scientific).

### Immunofluorescence

ARPE-19 cells were fixed with 4% (w/v) paraformaldehyde in PBS for 10 minutes at room temperature (RT) and incubated in blocking buffer (5% (w/v) bovine serum albumin (BSA, A3059, Sigma-Aldrich) and 0.5% Triton X-100 (A16046, Thermo Fisher Scientific) in PBS) at RT for 1 hour, followed by incubation at 4 °C overnight with the appropriate primary antibody (diluted in blocking buffer). The cells were then rinsed with PBS three times and incubated with the corresponding secondary antibody (diluted in blocking buffer) at RT for 1 hour. Upon washing with PBS three times, the sample was stained with 1 µg/mL DAPI (MBD0015, Sigma-Aldrich) in PBS for 10 minutes at RT, and washed again in PBS for two times prior to imaging.

### Western blotting

Total protein was extracted from cultured cells by lysis in RIPA buffer (R0278, Sigma-Aldrich) and sonicated briefly on a Vibra-Cell ultrasonic processor (VC-505, Sonics and Materials) with 1 second pulses for a total of 10 seconds. The lysate was centrifuged, its protein concentration was measured with Pierce BCA Protein Assay Kit (23227, Thermo Fisher Scientific), and then loaded onto either a 4-15% Mini-PROTEAN TGX precast gel (4561084, Bio-Rad) together with a PageRuler or PageRuler Plus protein marker (26616 or 26619, Thermo Fisher Scientific), and separated by gel electrophoresis at 120 V for 30 min. Proteins were then transferred onto a Polyvinylidenedifluoride (PVDF) membrane via semi-dry transfer on a Trans-Blot Turbo Transfer System (1704150, Bio-Rad), and incubated with blocking solution (3% (w/v) nonfat dry milk powder (1706404, Bio-Rad) in 1× tris-buffered saline (BUF-3030-20X4L, 1st Base) with 0.1% Tween 20 (T2008687, Sinopharm) (TBST)) for 1 hour. The membrane was then incubated with the appropriate primary antibody diluted in blocking solution at 4 °C overnight, washed with TBST for three times, and incubated with the corresponding secondary antibody diluted in blocking solution at RT for 1 hour. The membrane was then washed with TBST for three times, and incubated with Clarity Western ECL Substrate (1705061, Bio-Rad) for 10 minutes. The membranes were then visualized using an ImageQuant LAS 4000 digital imaging system (GE Healthcare Life Sciences).

### Antibodies

The following antibodies were used for immunofluorescence and Western blot experiments in this study: mouse anti-p21 (MA5-31479, Invitrogen), rabbit anti-p21 (MA5-14949, Invitrogen), rabbit anti-phospho-Rb (Ser807/811) (8516, Cell Signaling Technology), mouse anti-lamin A/C (4777, Cell Signaling Technology), mouse anti-lamin A (ab8980, Abcam), rabbit anti-lamin B1 (ab16048, Abcam), rabbit anti-LBR (ab32535, Abcam), mouse anti-LBR (ab232731, Abcam), rabbit anti-TMPO (*i.e*. LAP2β) (HPA070805, Sigma-Aldrich), mouse anti-emerin (ab204987, Abcam), rabbit anti-emerin (30853, Cell Signaling Technology), mouse anti-GAPDH (MA5-15738, Invitrogen), mouse anti-HaloTag (G9211, Promega), rabbit anti-HP1 alpha (ab109028, Abcam), Alexa Fluor 488-labeled goat anti-mouse IgG H&L (ab150113, Abcam), Alexa Fluor 647-labeled goat anti-rabbit IgG H&L (ab150079, Abcam), HRP-linked goat anti-rabbit IgG H&L (7074, Cell Signaling Technology), and HRP-linked horse anti-mouse IgG H&L (7076, Cell Signaling Technology).

### RNA sequencing and bioinformatic analyses

Total RNAs were extracted from ARPE-19 cells without or with *t*BHP treatment (in three replicates each), and purified using PureLink RNA Mini Kit (12183025, Invitrogen) according to manufacturer’s protocol. RNA sample purity and integrity were examined with a NanoDrop 2000 spectrophotometer (Thermo Fisher Scientific), and submitted for whole-transcriptome sequencing and analysis by BGI Genomics (Shenzhen, China). Briefly, sequencing was run on a DNBSEQ platform with paired-end reads of 150 base pairs, generating an average of 33 million clean reads per sample that were mapped to reference human genome (GRCh38) with 98.6% alignment rate, detecting a total of 16,486 genes. Sequencing readouts, alignments, and analysis results (including KEGG pathway enrichment and GO cellular component enrichment analyses) were retrieved from Dr. Tom (https://www.bgi.com/global/service/dr-tom), an interactive data analysis platform provided by BGI Genomics. Comparison with existing LADs database in RPE-1 cells (56) (retrieved from the 4D Nucleome Data Portal (https://data.4dnucleome.org/; 58, 59) under accession number 4DNESENITEIC) was performed using the Integrative Genomics Viewer tool (60).

### CRISPR cell lines generation

To generate CRISPR/Cas9 knock-in ARPE-19 cell lines that stably expresses either LBR-Halo or Halo-emerin at endogenous level, sgRNAs targeting near the stop codon of LBR gene or the transcription start site of EMD gene were designed using the Horizon Discovery CRISPR Design Tool online (https://horizondiscovery.com/en/ordering-and-calculation-tools/crispr-design-tool). The respective sgRNA was cloned into pCAG-spCas9-mCherry vector (a gift of Tan Meng How, Nanyang Technological University) by Gibson assembly upon digestion with BbsI-HF restriction enzyme (R3539S, New England Biolabs). To generate cell line stably expressing LBR-Halo, the homology-directed repair (HDR) template DNA sequence (with an added HaloTag sequence) that flanks the cut site near the stop codon of LBR gene was designed in-house and synthesized as a gBlocks gene fragment (Integrated DNA Technologies), and further amplified by polymerase chain reaction (PCR) to yield sufficient quantity for transfection. To generate cell line stably expressing Halo-emerin, HDR template DNA sequence flanking the cut site near the start codon of EMD gene was generated from the ARPE-19 cell genome by PCR, and ligated onto an empty PCR-linearized pCS2+ vector together with a HaloTag sequence via a series of Gibson assembly reactions using the NEBuilder HIFI DNA Assembly Master Mix (E2621S, New England Biolabs). PCR or Gibson assembly product for either donor template sequence was purified and co-transfected with CRISPR/Cas9 vector into ARPE-19 cells using Lipofectamine 3000 (L3000-015, Thermo Fisher Scientific); positive cells were enriched by FACS (detected at 561 nm, upon labeling with Janelia Fluor 549 (JF_549_) dye via HaloTag), and further validated by microscopy and Western blotting (**Fig. S3**). The sgRNAs, HDR templates and primers used are listed in **Tables S1** to **S3**; all constructs generated were validated by sequencing before transfection.

### Live-cell labelling

ARPE-19 cells expressing LBR-Halo or Halo-emerin were labelled in supplemented DMEM/F12 medium containing 5 nM JF_549_ HaloTag ligand (GA1110, Promega) at 37 °C for 30 minutes in a humidified incubator under 5% CO_2_, while co-staining the cell nucleus with NucBlue Live ReadyProbes Reagent (Hoechst 33342, R37605, Invitrogen). Cells were then gently rinsed with PBS once, and washed three times with fresh supplemented DMEM/F12 medium at 37 °C for 10 minutes each to remove the excess unlabeled ligand.

### Microscope setup

All live-cell imaging FCS and SMT measurements were performed on a customized Eclipse Ti2-E inverted microscope (Nikon) equipped for highly inclined and laminated optical sheet (HILO) illumination (40), with 4 laser lines at excitation wavelengths of 405 nm (60 mW, LuxX 405-60, Omicron), 488 nm (200 mW, LuxX 488-200, Omicron), 561 nm (150 mW, OBIS 1280720, Coherent) and 647 nm (200 mW, LuxX 638-200, Omicron), respectively. Live-cell samples were imaged at 37 °C under a humidified 5% CO_2_ atmosphere using a 100x, NA 1.49 oil objective (SR HP Apo, Nikon). Images were acquired using the NIS-Elements AR software (version 5.21.03, Nikon), and recorded with an EMCCD camera (Andor iXon Life 897, Oxford Instruments).

### Confocal microscopy

Immunofluorescently labelled cells were imaged on a Nikon Eclipse Ti inverted microscope equipped with a Nikon C2si confocal scanhead and 4 laser lines at excitation wavelengths of 405 nm (100 mW, OBIS 1284371, Coherent), 488 nm (150 mW, OBIS 1277611, Coherent), 561 nm (150 mW, OBIS 1280720, Coherent) and 639 nm (MRL-FN-639-300mW, Changchun New Industries Optoelectronics Tech. Co.), respectively. Confocal images were acquired with a 60x, NA 1.20 water immersion objective (CFI Plan Apochromat VC 60XC WI, Nikon) or a 100x, NA 1.45 oil immersion objective (CFI Plan Apochromat Lambda D 100X Oil, Nikon), using the NIS-Element AR software (version 5.21.02, Nikon).

### Imaging FCS and SMT measurements

For imaging FCS measurements, live ARPE-19 cells expressing LBR-Halo or Halo-emerin (labelled by JF_549_ HaloTag ligand) were illuminated at 561 nm under HILO mode at a power density of ∼0.02 kW/cm^2^ and an acquisition speed of 5.5 ms per frame, for up to 40,000 frames with a ROI of 128 × 128 pixels. For SMT measurements, live ARPE-19 cells expressing LBR-Halo or Halo-emerin (labelled by JF_549_ HaloTag ligand) were illuminated at 561 nm under HILO mode at a power density of ∼0.02 kW/cm^2^ and an acquisition speed of 300 ms per frame, for up to 1,000 frames with a ROI of 256 × 256 pixels. Hoechst 33342 images of each cell nucleus were acquired before and after each measurement to be used as a mask for isolating intranuclear trajectories in subsequent image analysis. For correlative mapping experiments, cells were immediately fixed upon imaging FCS measurements with 4% (w/v) paraformaldehyde in PBS for 10 minutes at RT, before proceeding with immunofluorescence labeling.

### Imaging FCS data analysis

All image processing/analyses for imaging FCS/SMT/STAR mapping, numerical simulations, and data visualization were performed with MATLAB (MathWorks) using custom-written routines described below. To analyze the imaging FCS data, the fluorescence intensity trace recorded at each pixel of an image sequence was extracted over time, and corrected for photobleaching by fitting and removing the slowly varying intensity decay using a high-degree polynomial. The corrected intensity trace was then used to calculate the temporal autocorrelation function *G*(τ), which was fitted with a two-dimensional diffusion model to obtain local estimates of the diffusion coefficient *D* and molecular number *N*:

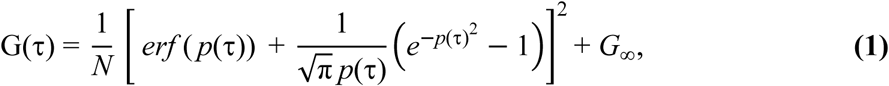

where 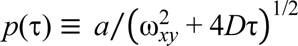, with *a* being the pixel size, ω*_xy_* being the lateral 1/*e*^2^ radius of the point-spread function, and *G*_∞_ being the autocorrelation offset at long lag times (set to zero in our analysis).

### SMT data analysis

SMT data analysis was adapted from previously described procedures (46) with modifications. Slow-tracking movies were acquired at an exposure time of 300 ms per frame, resulting in motion-blurring of rapidly diffusing molecules while slowly diffusing or bound molecules remained detectable as distinct bright spots. Individual frames were first denoised using a wavelet filter (61), and the two-dimensional coordinates of single-molecule localizations were determined using a radial gradient-based localization algorithm (62). Trajectories were reconstructed from the resulting localizations using the TrackMate algorithm (63). Tracks were required to contain at least 3 detections (*i.e.* 2 consecutive displacements). The maximum displacement between adjacent frames was set to 280 nm, and up to 2 consecutive missed detections were allowed during trajectory linking. Binding events were identified from reconstructed trajectories based on their spatial confinement. To do so, all possible sub-trajectories were evaluated, and their spatial extent (*R_g_*) was estimated from the spatial distribution of their localization coordinates. A sub-trajectory was classified as a binding event when its spatial extent remained below a duration-dependent confinement threshold:

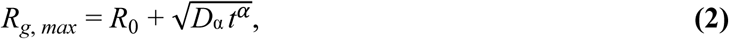

where *t* is the duration of the sub-trajectory, *R*_0_ is the offset introduced by measurement error, *D*_α_ is the apparent diffusion coefficient of the bound molecule, and α is the corresponding anomalous diffusion coefficient. Here, a duration-dependent confinement threshold was used to account for the small spatial displacements that can occur even while a molecule remains bound, leading to a progressively larger spatial extent for longer binding events. The values for *R*_0_ (40 nm), *D*_α_ (0.0016 μm^2^s^−α^) and α (0.5) were empirically set to provide an appropriate confinement criterion across binding durations, and are consistent with the heterogeneous and highly confined dynamics observed for inner nuclear membrane proteins such as LBR and emerin (64, 65). For each trajectory, the longest sub-trajectory satisfying the confinement criterion was first selected, after which the remaining portions were iteratively evaluated to identify additional binding events. Only binding events containing at least 4 consecutive displacements were retained, thereby minimizing the contribution of slowly diffusing molecules that could transiently resemble bound molecules. The duration of each identified binding event was used to construct the survival probability, *S*(*t*), defined as the probability that a molecule remains bound for at least a time *t*. Survival probability curves were fitted with a triple-exponential model:

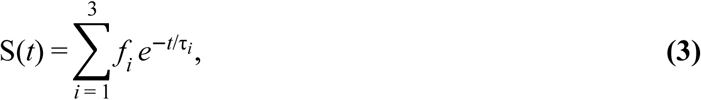

where *t_i_* is the characteristic residence time and *f_i_* the corresponding molecular fraction of each binding mode, with the three fractions summing to 1. Binding events were classified as transient or stable according to their duration, with events shorter than 1.8 s classified as transient and those lasting at least 1.8 s classified as stable (*i.e.* corresponding to 6 consecutive displacements at the 300-ms acquisition time).

### Spatial analysis of binding events

To perform STAR mapping of binding events, a reconstruction grid with a pixel size of 40 nm was defined, and the midpoint of each consecutive displacement in a trajectory was assigned to the corresponding grid pixel. The intensity of each pixel therefore reflected the number of displacements falling within that pixel. The resulting displacement-density maps were smoothened using a Gaussian filter (σ = 40 nm) and independently rescaled for visualization, with transient and stable binding events displayed as separate channels in composite maps. Stable binding events were further analyzed for spatial clustering, in which each event was represented by its mean spatial position and pairwise distances between them were calculated. Events separated by no more than 200 nm were considered spatially clustered, and each cluster containing at least 3 stable binding events was defined as a binding hotspot. Stable events not belonging to such clusters were classified as unclustered. Displacement-density maps of clustered and unclustered stable binding events were generated using the same 40-nm grid and Gaussian smoothening (σ = 40 nm) and displayed as separate channels.

To quantitatively characterize the spatial features of binding events, the following parameters were quantified from the above analysis: Targeting efficiency is defined as the ratio between the NL area sampled by stable binding and that by diffusion and transient binding combined; stable binding density or hotspot density is defined as the number of stable binding events or binding hotspots per unit area of NL; extent of clustering is defined as the ratio between the numbers of stable binding events clustered within the hotspots and those outside the hotspots (*i.e.* unclustered); lastly, the lifetime for each hotspot is defined as the time interval between the first and last stable binding events within that hotspot.

### Correlative radial profile analysis

To quantify the spatial relationship between HP1α organization and LBR mobility, radial profiles of HP1α fluorescence intensity and LBR diffusion coefficients were calculated around individual HP1α foci. Firstly, HP1α distribution at the NL was intensity-normalized between 0 and 1, and segmented by watershed analysis into distinct foci. To limit over-segmentation of individual HP1α foci, weak local minima separated from neighboring minima by intensity barriers smaller than 0.2 were suppressed before watershed segmentation. Segmentation was restricted to regions within the selected ROI corresponding to the basal surface of the NL with normalized HP1α intensity > 0.5, while foci with centroid position within 20 pixels (800 nm) from the boundary of the ROI were excluded to ensure that the entire radial profile remains within the ROI. A minimum area of 10 pixels² (0.016 µm²) was used as cutoff for all foci, whereas a minimum area of 250 pixels² (0.4 µm²) was used as cutoff for large foci corresponding to compacted heterochromatin. All segmentation parameters were kept constant across the experimental conditions. For each focus thus identified, HP1α fluorescence intensity and LBR diffusion coefficients were sampled radially from its intensity-weighted centroid over a distance of 800 nm (*i.e.* 20 pixels for a 40 nm/pixel grid) along 200 uniformly distributed angular directions. At each radial distance, values were averaged over all directions to generate one-dimensional radial profiles of HP1α intensity and LBR diffusion coefficient. To quantify changes in LBR mobility across each HP1α focus, the mean LBR diffusion coefficient for the inner region of the focus (defined as the first five radial points, *i.e.* 0– 160 nm) was compared with the mean diffusion coefficient for the outer region (defined as the last five radial points, *i.e.* 640–800 nm); the relative radial change in LBR diffusion coefficient was then calculated as:

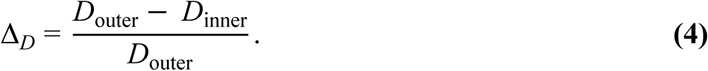

Foci with Δ*_D_* > 0.2 were classified as exhibiting decreased LBR mobility toward the center of HP1α foci (*i.e.* anti-correlated), whereas domains with Δ*_D_* < −0.2 were classified as exhibiting increased LBR mobility toward the center (*i.e.* correlated). Foci that show no substantial radial change (*i.e.* with −0.2 ≤ Δ*_D_* ≤ 0.2) were classified as non-correlated.

### Numerical simulations

To establish a null reference model for the radial profile analysis described above, synthetic HP1α distribution and LBR mobility maps were numerically simulated. These simulations were not intended to provide a biophysical model for the mechanistic underpinnings of the experimentally observed spatial patterns; rather, they provided artificial datasets with realistic statistical and spatial properties to which the same radial profile analysis used for the experimental data could be applied. They therefore established the level of apparent radial association expected in the absence of an underlying spatial relationship between the two signals. Synthetic HP1α maps were generated by randomly positioning Gaussian intensity domains, with their amplitudes and widths jointly sampled from the experimentally measured amplitude–width distribution and individual Gaussian widths restricted to σ ≤ 4 pixels. Synthetic LBR mobility maps were generated by combining two independent Gaussian random fields smoothened with Gaussian kernels of σ = 1 and 2 pixels, respectively. These two spatial scales were chosen to introduce shorter- and longer-range variations qualitatively resembling the spatial texture of the experimental mobility maps, rather than to quantitatively reproduce their spatial correlation structure. The resulting field was then histogram-matched to the experimental diffusion-coefficient distribution using values sampled from a beta distribution fitted to the experimental data. This procedure reproduced the experimental distribution of diffusion coefficients while retaining a qualitatively realistic degree of spatial heterogeneity.

### Statistical analysis

All data presented were derived from at least three independent measurements or replicates. For line plots and histograms, the mean and standard deviation are shown. For box-and-whisker plots, the median, 25^th^/75^th^ percentiles (box) and 5^th^/95^th^ percentiles (whiskers) are shown. Statistical significance was assessed using a two-tailed unpaired Student’s *t*-test or a two-tailed Welch’s *t*-test (in **Fig. 6**): *: *p* < 0.05; **: *p* < 0.01; ***: *p* < 0.001; NS: not significant.

## DATA AND CODE AVAILABILITY

RNA sequencing data on our RPE cellular senescence model will be deposited at NCBI Gene Expression Omnibus (GEO) and made publicly available prior to publication. All codes used in this study will be deposited at GitHub and made publicly available prior to publication. Due to their large size, the raw imaging FCS and SMT data are available from the corresponding authors upon reasonable request.

## AUTHOR CONTRIBUTIONS

Z.W.Z. conceived the study; Z.W.Z. and X.K. designed the experiments; X.K. performed all experiments; W.E. performed data analyses of imaging FCS, SMT and STAR/correlative mapping as well as numerical simulations; Z.W.Z. and P.K. co-supervised the study; Z.W.Z. and X.K. wrote the manuscript with contributions from all authors.

## ACKNOWLEDGEMENTS

We thank Meng How Tan for the gift of the pCAG-spCas9-mCherry plasmid, Xinyi Su for the gift of ARPE-19 cell line, Marie Cutiongco for the gift of lamin A antibody, and Rong Li and Thorsten Wohland for helpful discussions and suggestions. This work was supported by two Ministry of Education of Singapore Academic Research Fund Tier 2 Grants (MOE-T2EP30222-0010 and MOE-001004-00) to Z.W.Z. and one Tier 3 Grant (MOET32020-0001) to P.K. and Z.W.Z., as well as a National Research Foundation of Singapore Mid-Sized Grant (NRF-MSG-2023-0001) to P.K.

## Supporting Material for

### This PDF file includes

Figures S1 to S10

Legends for Movies S1 to S2

Tables S1 to S3

### Other supporting materials for this manuscript include

Movies S1 to S2

**Figure S1.**
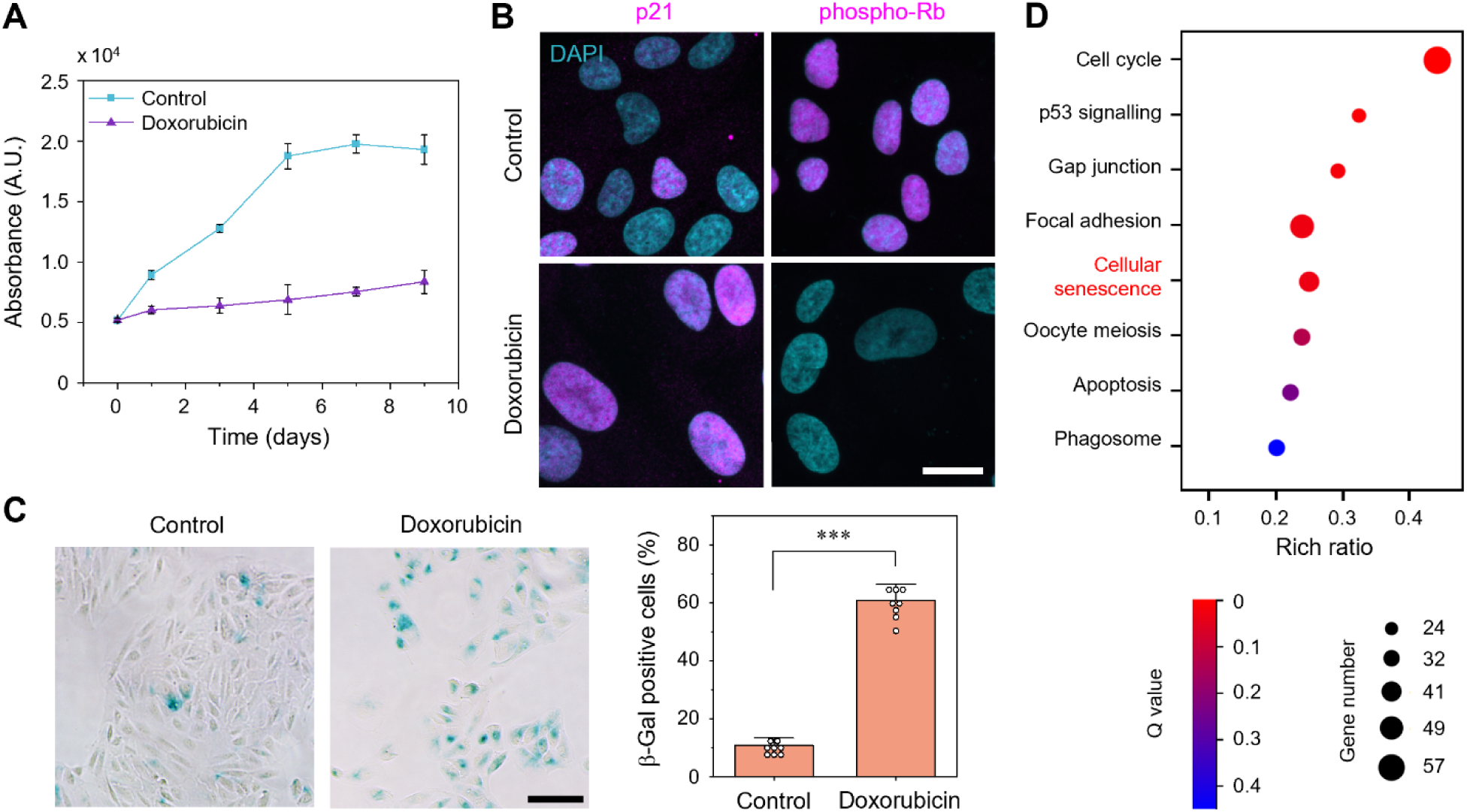
Characterization of senescence induction in ARPE-19 cells with doxorubicin treatment. **(A)** Cell proliferation profiles of doxorubicin-treated and untreated ARPE-19 cells (as measured by CCK-8 assay) over a period of 9 days; day 0 denotes the commencement of the first dose of doxorubicin treatment. **(B)** Representative confocal images of doxorubicin-treated and untreated ARPE-19 cells immunofluorescently stained for cell cycle arrest markers p21 or phospho-Rb; the cell nucleus is co-stained with DAPI. **(C)** Representative bright-field images of doxorubicin-treated and untreated ARPE-19 cells stained for senescence-associated β-galactosidase (left), as well as the percentage of β-galactosidase-positive cells in each sample (right). **(D)** Kegg pathway enrichment analysis of RNA-seq data on doxorubicin-treated and untreated ARPE-19 cells, showing the top cellular processes-related hits upon doxorubicin treatment (including cellular senescence). Scale bar: 20 µm (**B**); 100 µm (**C**). *n* = 3 (**A**) and 8 (**C**) independent replicates; data are represented as mean ± S.D..

**Figure S2.**
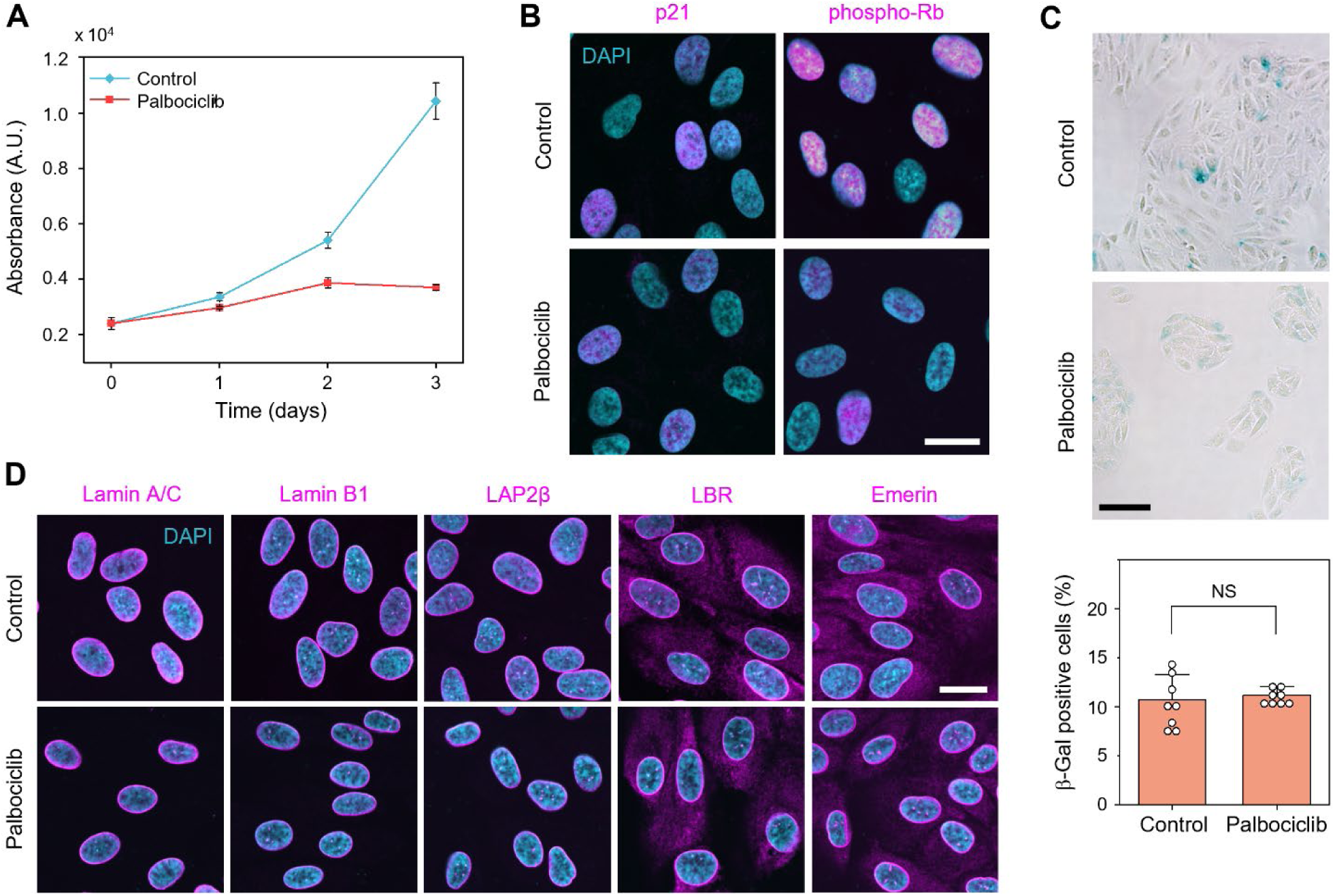
Senescence-induced expression-level changes for key NL components are not a consequence of non-senescence-associated cell cycle arrest. **(A)** Cell proliferation profiles of palbociclib-treated and untreated ARPE-19 cells (as measured by CCK-8 assay) over a period of 3 days; day 0 denotes the commencement of palbociclib treatment. Measurement terminates at day 3 as proliferation has stopped for palbociclib-treated ARPE-19 cells. **(B)** Representative confocal images of palbociclib-treated and untreated ARPE-19 cells immunofluorescently stained for cell cycle arrest markers p21 or phospho-Rb; the cell nucleus is co-stained with DAPI. While phospho-Rb expression is reduced (similar to that observed for *t*BHP and doxorubicin treatments), the fact that the p53–p21 pathway is not affected suggests that the cell cycle arrest caused by palbociclib treatment is likely not a consequence of senescence. **(C)** Representative bright-field images of palbociclib-treated and untreated ARPE-19 cells stained for senescence-associated β-galactosidase (top), as well as the percentage of β-galactosidase-positive cells in each sample (bottom), further indicating that palbociclib treatment does not induce senescence in ARPE-19 cells. The image for untreated ARPE-19 cells is the same as that in **Fig. S1C**. **(D)** Representative confocal images of palbociclib-treated and untreated ARPE-19 cells immunofluorescently stained for each of 6 key components of the NL: lamin A/C, lamin B1, LAP2β, LBR, and emerin; the cell nucleus is co-stained with DAPI. In particular, palbociclib treatment does not lead to significant reduction in the levels of lamin B1, LAP2β or LBR, as was observed for *t*BHP treatment. Scale bar: 20 µm (**B** and **D**); 100 µm (**C**). *n* = 3 (**A**) and 8 (**C**) independent replicates; data are represented as mean ± S.D..

**Figure S3.**
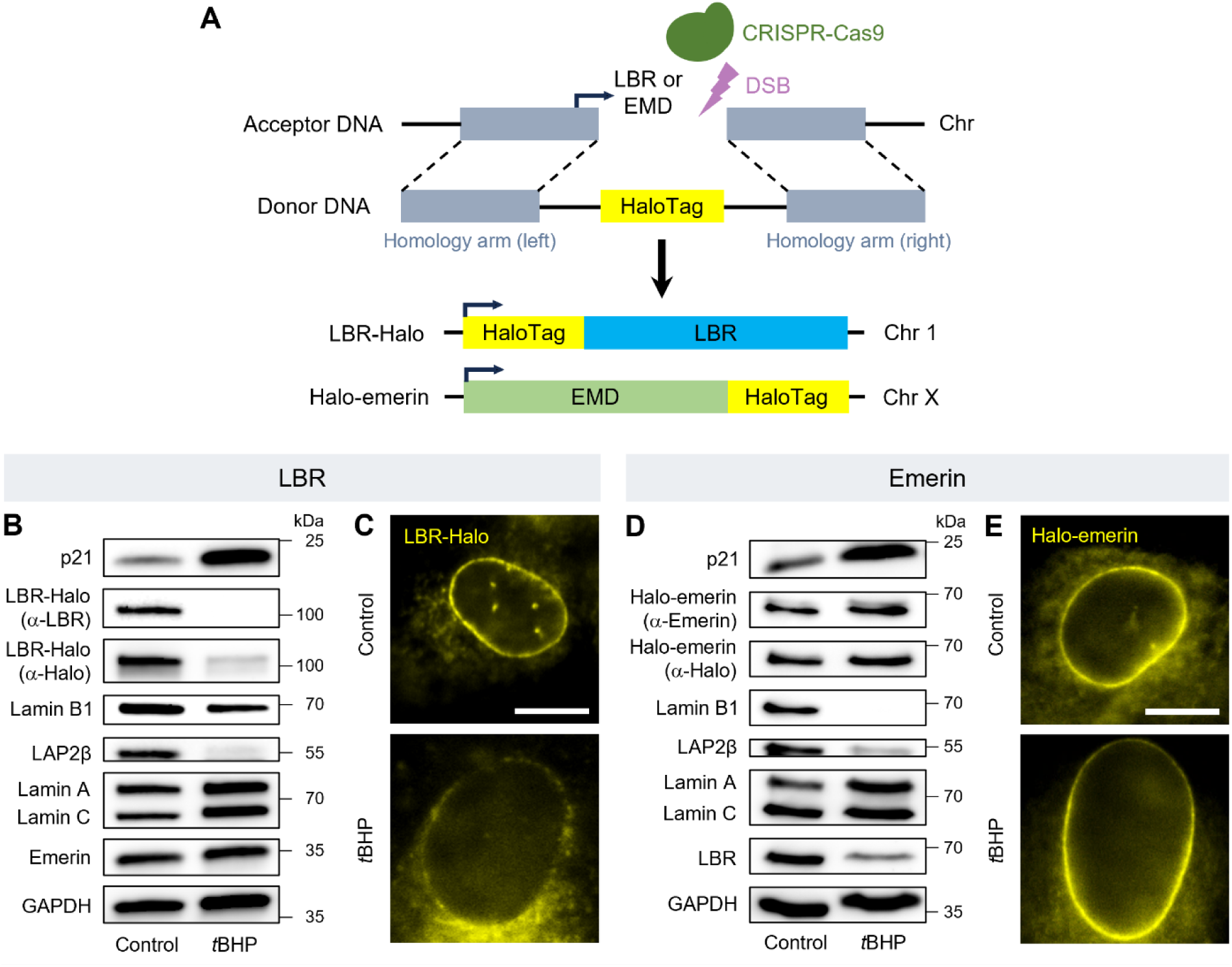
Design and validation of CRISPR knock-in ARPE-19 cell lines endogenously expressing HaloTag-fused LBR or emerin. **(A)** Schematic for the generation of knock-in ARPE-19 cell lines expressing LBR-Halo or Halo-emerin at endogenous level, using CRISPR/Cas9-mediated genome editing. **(B** and **D)** Western blots for LBR-Halo **(B)** or Halo-emerin **(D)**, together with p21 and the other 5 key NL components in *t*BHP-treated and untreated CRISPR knock-in cells; GAPDH serves as loading control. The similar patterns of expression-level changes as those observed for wild-type ARPE-19 cells indicate that our knock-in cell lines can be similarly induced into a senescent state with *t*BHP treatment. **(C** and **E)** Representative images of live *t*BHP-treated and untreated CRISPR knock-in cells expressing LBR-Halo **(C)** or Halo-emerin **(E)** (each labeled with JF_549_ dye), acquired under HILO illumination with similar imaging conditions as SMT measurements. The substantial nuclear size enlargement as well as the distinct alteration patterns for the levels of LBR and emerin at the NL of *t*BHP-treated cells again indicate successful induction into a senescent state (The reduction in LBR-Halo signal from the nuclear periphery upon *t*BHP treatment appears less drastic as compared to that observed in Western blot **(B)** or immunofluorescence imaging (Fig. 2B) due to the much higher sensitivity of single-molecule imaging used here). Scale bar for **C** and **E**: 10 µm.

**Figure S4.**
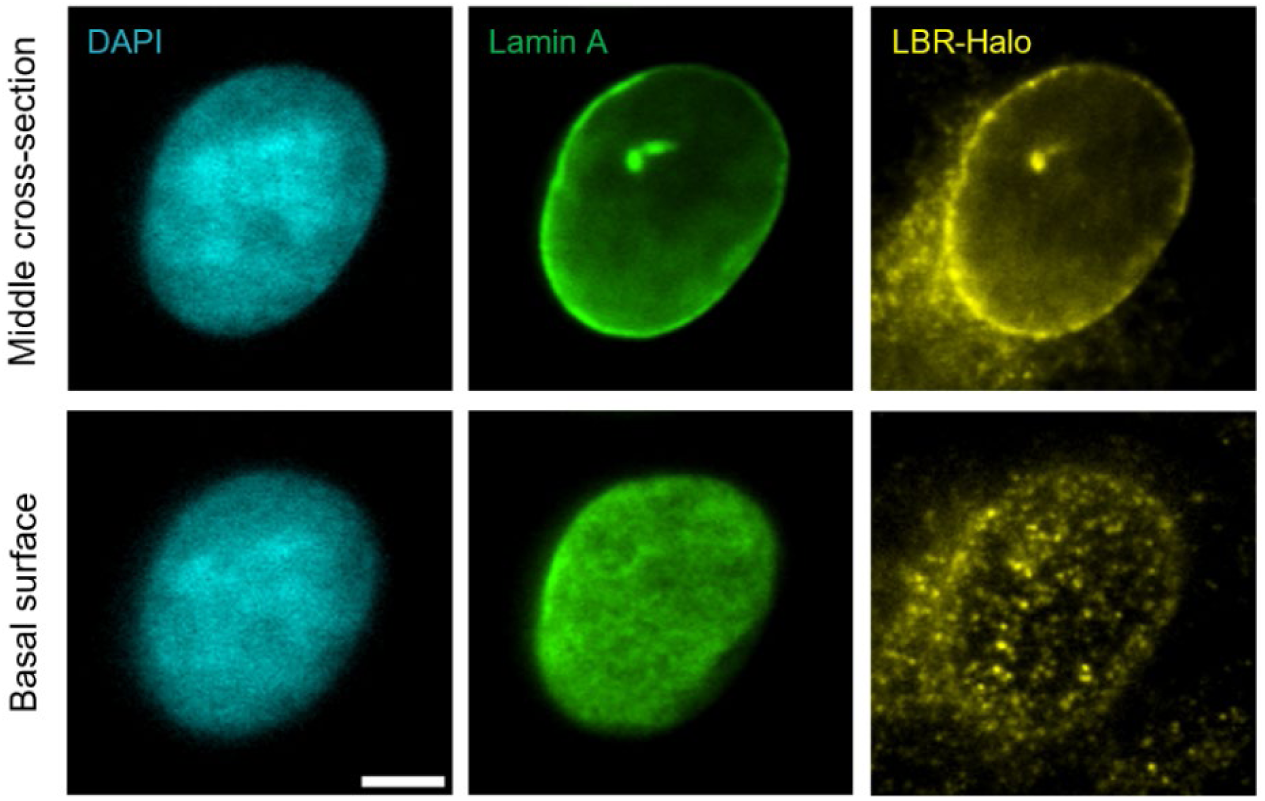
Acquiring imaging FCS and SMT measurements at the basal surface of the nuclear envelope. Representative HILO images of JF_549_-labeled LBR-Halo at either the middle cross-section of the cell nucleus (top row) or basal surface of the nuclear envelope (bottom row) in knock-in ARPE-19 cells, acquired under similar imaging conditions as SMT measurements; the cell nucleus is co-stained with DAPI, while lamin A (immunofluorescently labeled) serves as a positional marker for the NL. The distinctly different and much stronger LBR-Halo signal obtainable across the basal surface of the nuclear envelope (and detectable at single-molecule sensitivity) enables us to position the HILO illumination at precisely the basal surface of the nuclear envelope for the acquisition of imaging FCS and SMT measurements. Scale bar: 5 µm.

**Figure S5.**
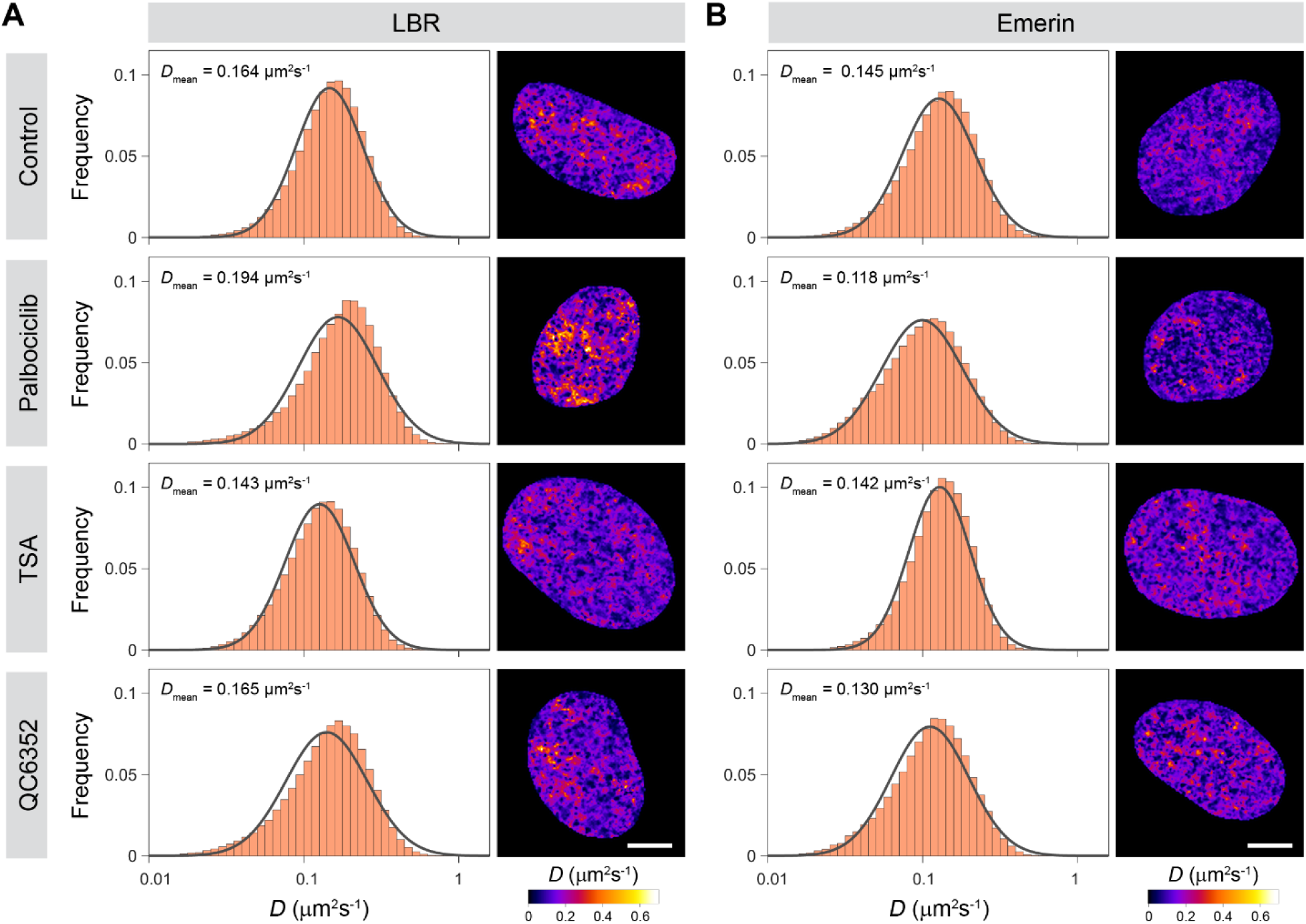
The mobilities of LBR and emerin are minimally impacted by palbociclib, TSA or QC6352 treatment. Histograms of diffusion coefficient (*D*, left column) as well as the corresponding mobility maps (right column) quantified from imaging FCS measurements on knock-in ARPE-19 cells endogenously expressing LBR-Halo **(A)** or Halo-emerin **(B),** without or with palbociclib, TSA or QC6352 treatment. Black curves in the histograms denote single-Gaussian fits. The histograms and mobility maps for the control cells are reproduced from **Fig. 3B** and **C** for ease of comparison. Scale bar: 5 µm. *n* = 63 (control), 10 (palbociclib-treated), 29 (TSA-treated) and 28 (QC6352-treated) cells for **A**, and 47 (control), 10 (palbociclib-treated), 20 (TSA-treated) and 9 (QC6352-treated) cells for **B**.

**Figure S6.**
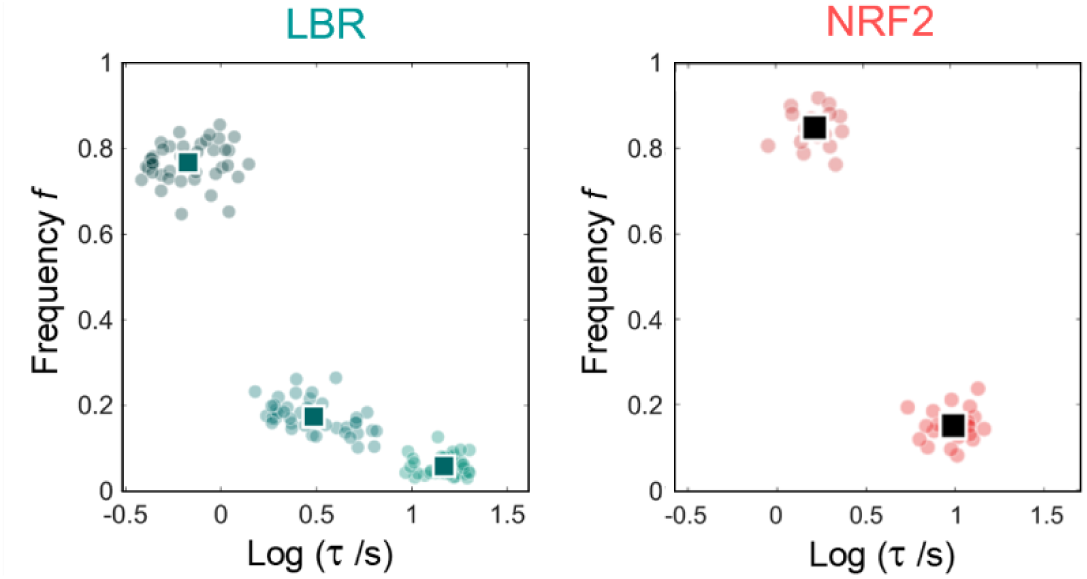
Validation for the 3-mode NL-binding dynamics observed. Scatter plots of residence time (τ) and frequency (*f*) associated with each binding mode resolved from live-cell SMT measurements on LBR (left, reproduced from Fig. 4C) and NRF2 (right), a transcription factor involved in oxidative stress response but is known not to bind the NL. In contrast to the 3-mode NL-binding dynamics observed for LBR, NRF2 exhibits only 2 binding modes with residence times distinctly different from those of LBR, hence validating the accuracy of our SMT measurements and analysis. Each dot denotes a single cell, and squares denote mean values. *n* = 43 (LBR) and 21 (NRF2) cells.

**Figure S7.**
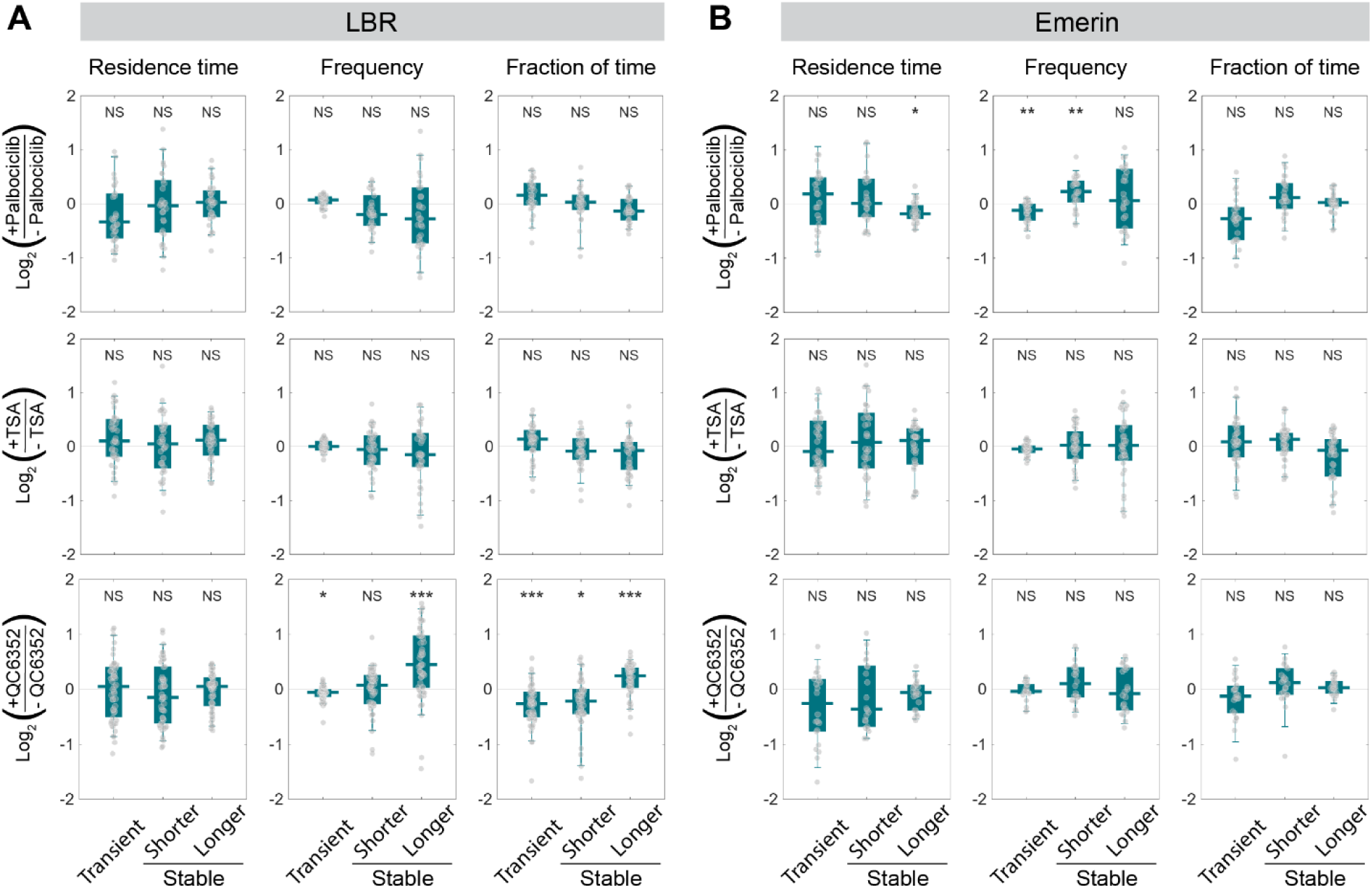
Alterations in NL-binding dynamics for LBR and emerin are not a consequence of non-senescence-associated cell cycle arrest or global changes in chromatin accessibility. Box-and-whisker plots of fold changes (on log_2_ scale) in residence time, binding frequency and fraction of time bound associated with each mode of NL-binding for LBR **(A)** and emerin **(B)**, upon treatment with palbociclib, TSA or QC6352. In contrast to the alterations observed for *t*BHP treatment, palbociclib treatment only moderately impacts the binding frequency of emerin without affecting the binding dynamics of LBR, while TSA treatment does not impact the binding dynamics of both. QC6352 treatment also does not impact the binding dynamics of emerin, but intriguingly shifts the molecular partitioning of LBR towards longer-lived stable binding (*i.e.* opposite of the trend observed for *t*BHP treatment), suggesting a potential role played by localized chromatin decondensation in modulating the NL-binding dynamics of LBR. Each dot denotes a single cell, with *n* = 94 (control), 30 (palbociclib-treated), 37 (TSA-treated) and 51 (QC6352-treated) cells for **A**, and 69 (control), 26 (palbociclib-treated), 36 (TSA-treated) and 22 (QC6352-treated) cells for **B**. Data are represented as median, 25^th^/75^th^ percentiles (box) and 5^th^/95^th^ percentiles (whiskers).

**Figure S8.**
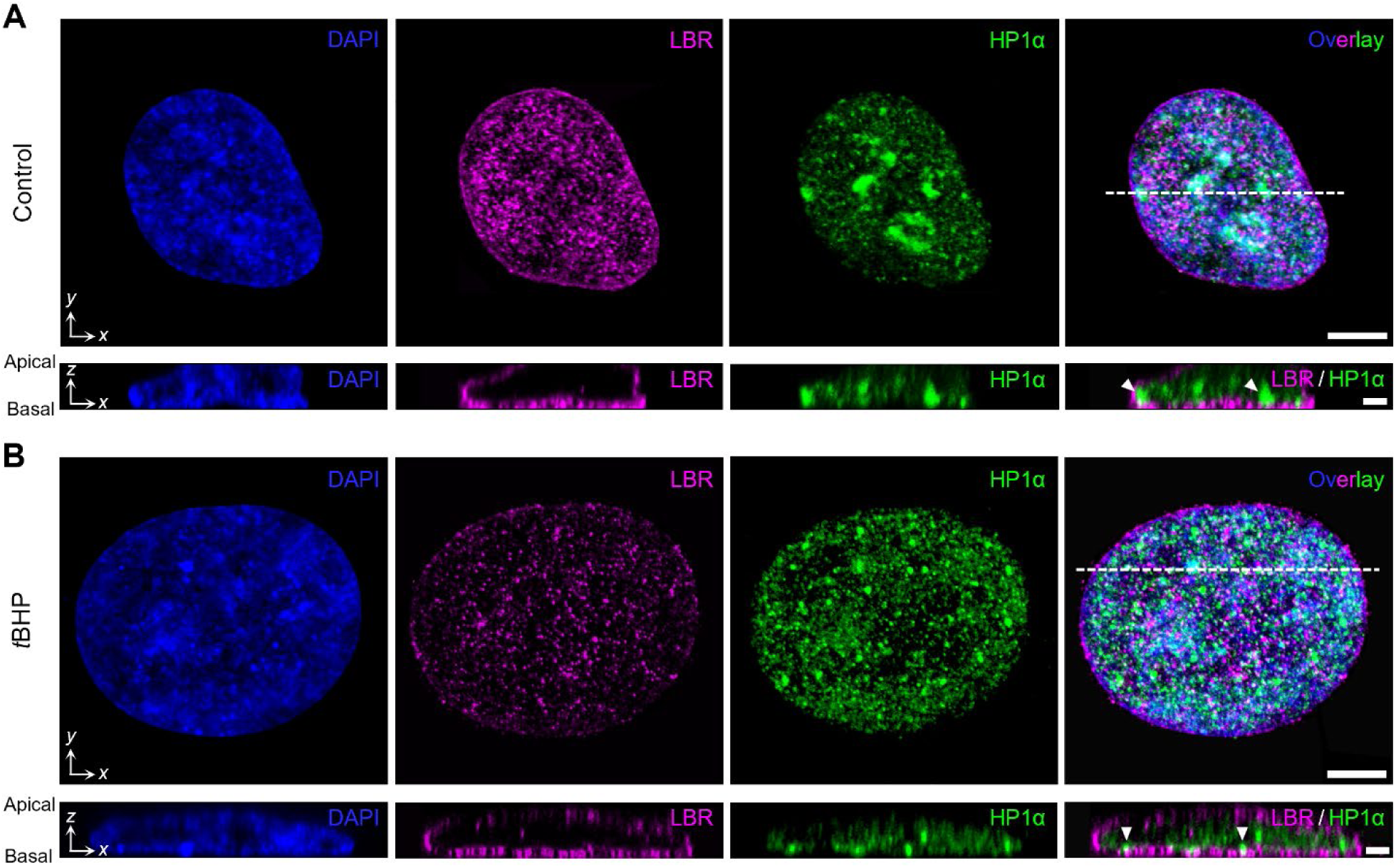
ARPE-19 cells undergo large-scale restructuring of constitutive heterochromatin domains during senescence. Representative confocal images of CRISPR knock-in ARPE-19 cells without **(A)** or with **(B)** *t*BHP treatment, showing intranuclear distributions of LBR and HP1α (both immunofluorescently labelled); the cell nucleus is co-stained with DAPI. Images in the top row are projections of a 3D confocal *z*-stack onto the *x*-*y* plane, while those in the bottom row are *x*-*z* profiles along the white dotted line in the overlay image of the *x*-*y* projections. White arrowheads indicate HP1α foci that are in contact with the basal surface of the nuclear lamina (as marked by LBR signal); the highly condensed large HP1α domains present in nonsenescent cells are dispersed into much smaller foci in *t*BHP-treated cells. Scale bar: 5 µm (*x*-*y* projections); 2 µm (*x*-*z* profiles).

**Figure S9.**
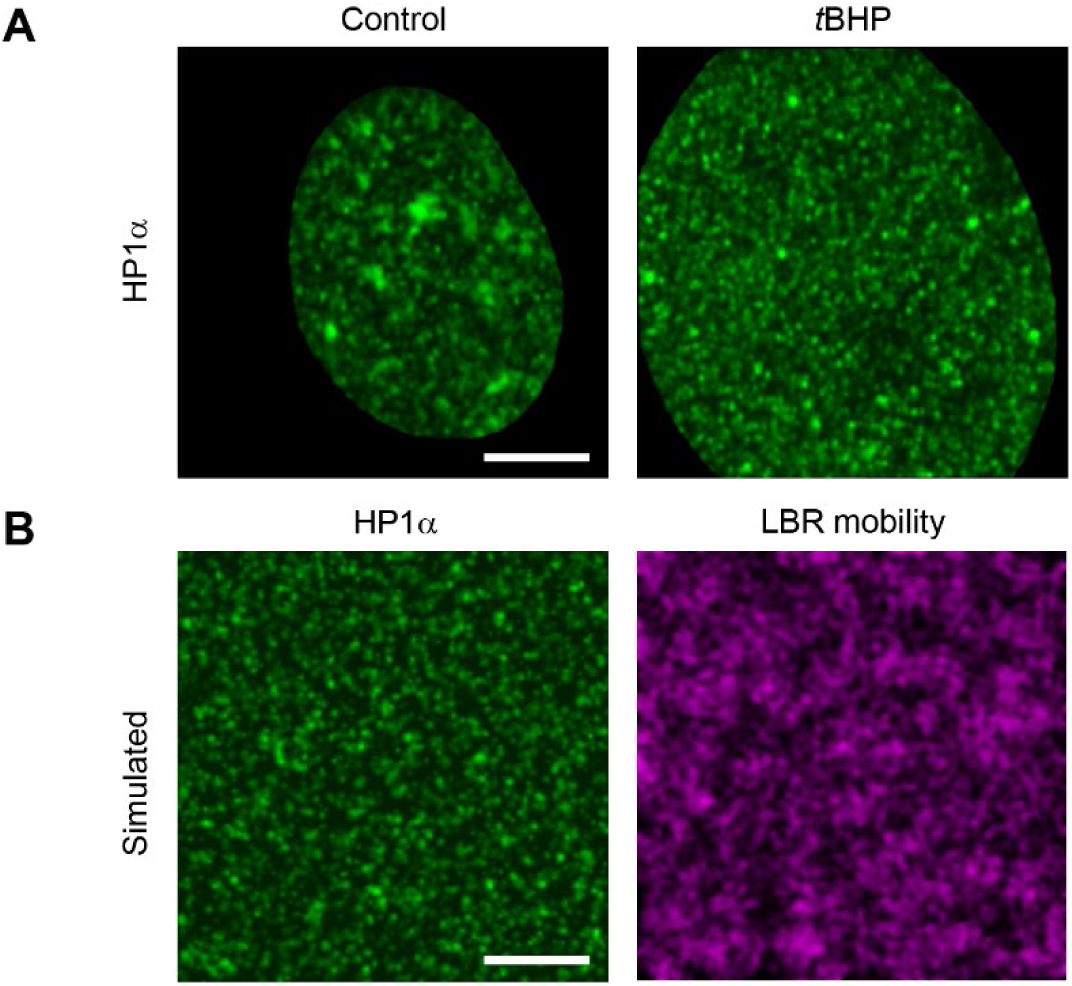
Comparison between experimentally observed and *in silico* simulated distributions of constitutive heterochromatin foci on the NL. **(A)** Unsegmented images of HP1α foci distribution on the NL basal surface in *t*BHP-untreated (left) and treated (right) ARPE-19 cells (corresponding to the segmented images in Fig. 6A). **(B)** Maps of HP1α foci (left) and LBR mobility (right) on the NL simulated *in silico* using relevant parameters derived from experimental imaging data (see **Materials and Methods** for details). The simulated distribution of HP1α foci highly resembles that observed in *t*BHP-treated senescent cells. Scale bar: 5 µm.

**Figure S10.**
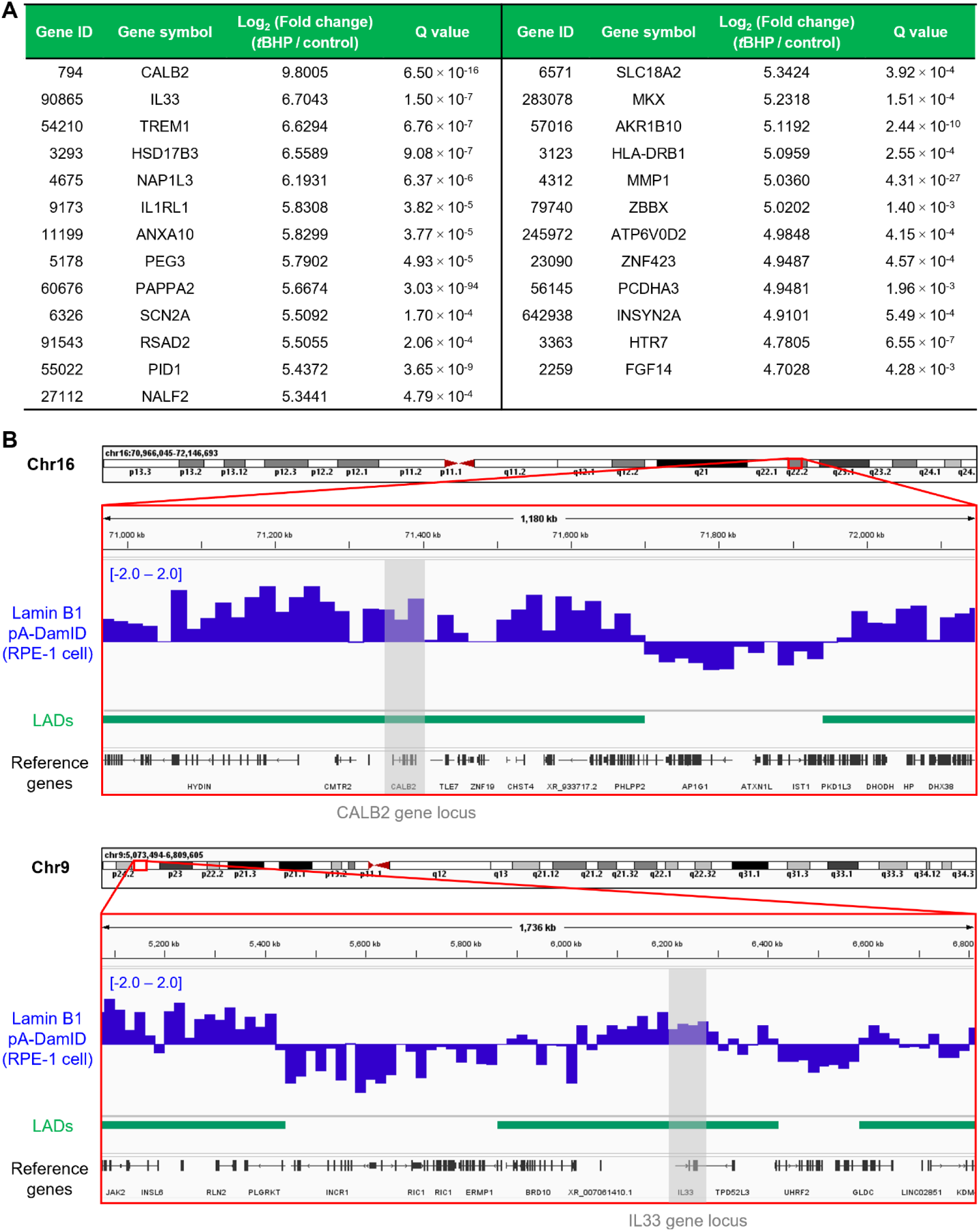
Upregulated genes during RPE cellular senescence exhibit significant association with the nuclear lamina. **(A)** A list of 25 genes (out of the 50 most upregulated genes detected in our *t*BHP-induced cellular senescence model) that fall within the genomic boundaries of known LADs, together with their associated fold changes and Q values. **(B)** Alignment of the genomic loci of the 2 most upregulated genes from **(A)** (CALB2 and IL33) against an existing database of LADs, profiled in RPE-1 cells with pA-DamID (56). The reference LADs database was retrieved from the 4D Nucleome Data Portal (https://data.4dnucleome.org/) under accession number 4DNESENITEIC.

**Movie S1: Representative dataset for imaging FCS.** A live ARPE-19 cell endogenously expressing LBR-Halo (and labeled with JF_549_-HaloTag ligand) is imaged under HILO illumination at 5.5 ms per frame. Scale bar: 2 µm.

**Movie S2: Representative dataset for SMT.** A live ARPE-19 cell endogenously expressing LBR-Halo (and labeled with JF_549_-HaloTag ligand) is imaged under HILO illumination at 300 ms per frame. Scale bar: 2 µm.

**Table S1:**
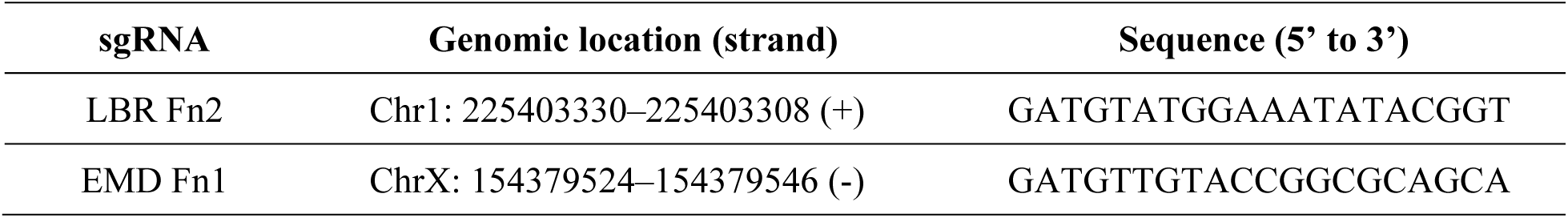
List of sgRNAs used for the generation of stable cell lines used in this study.

**Table S2:**
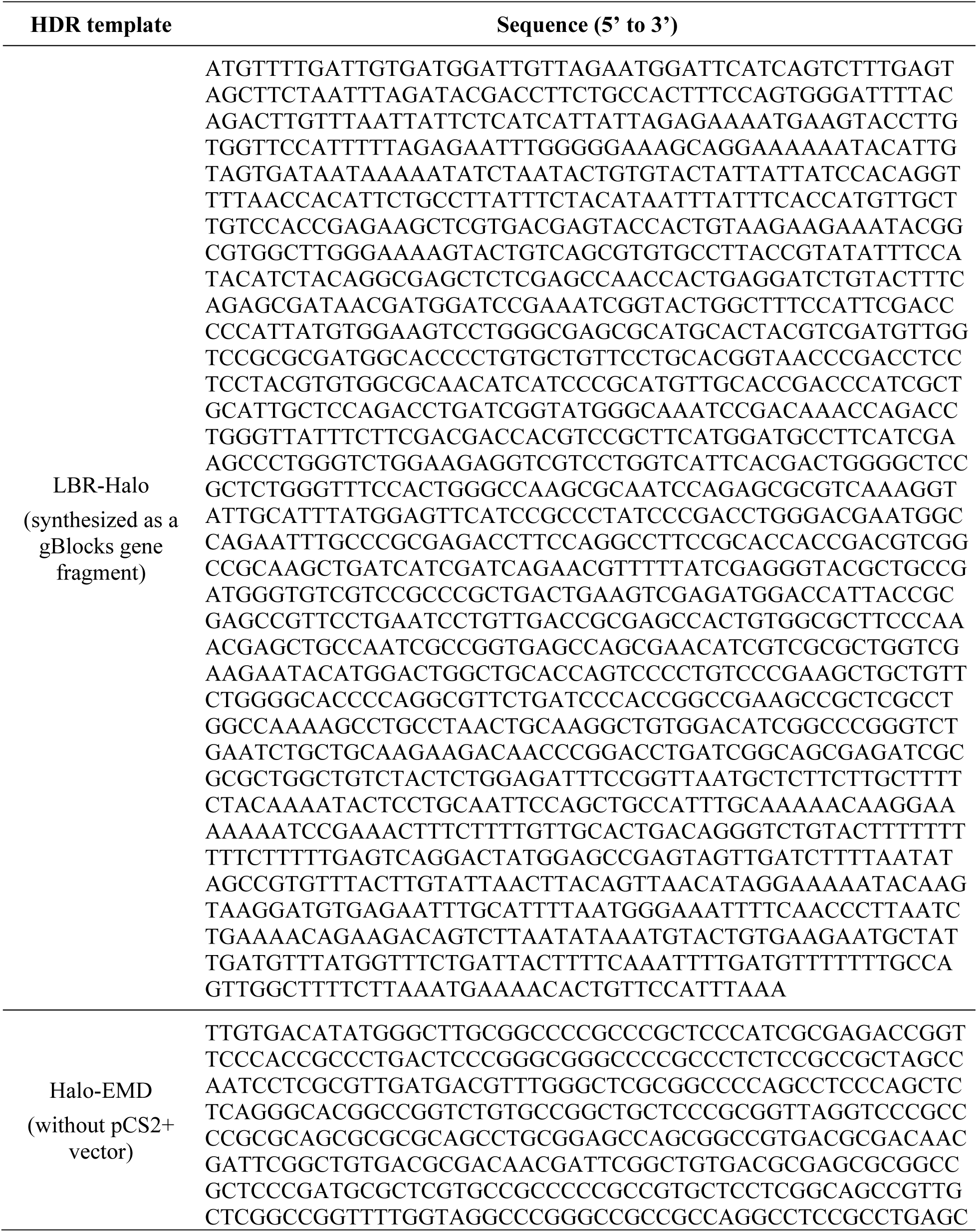
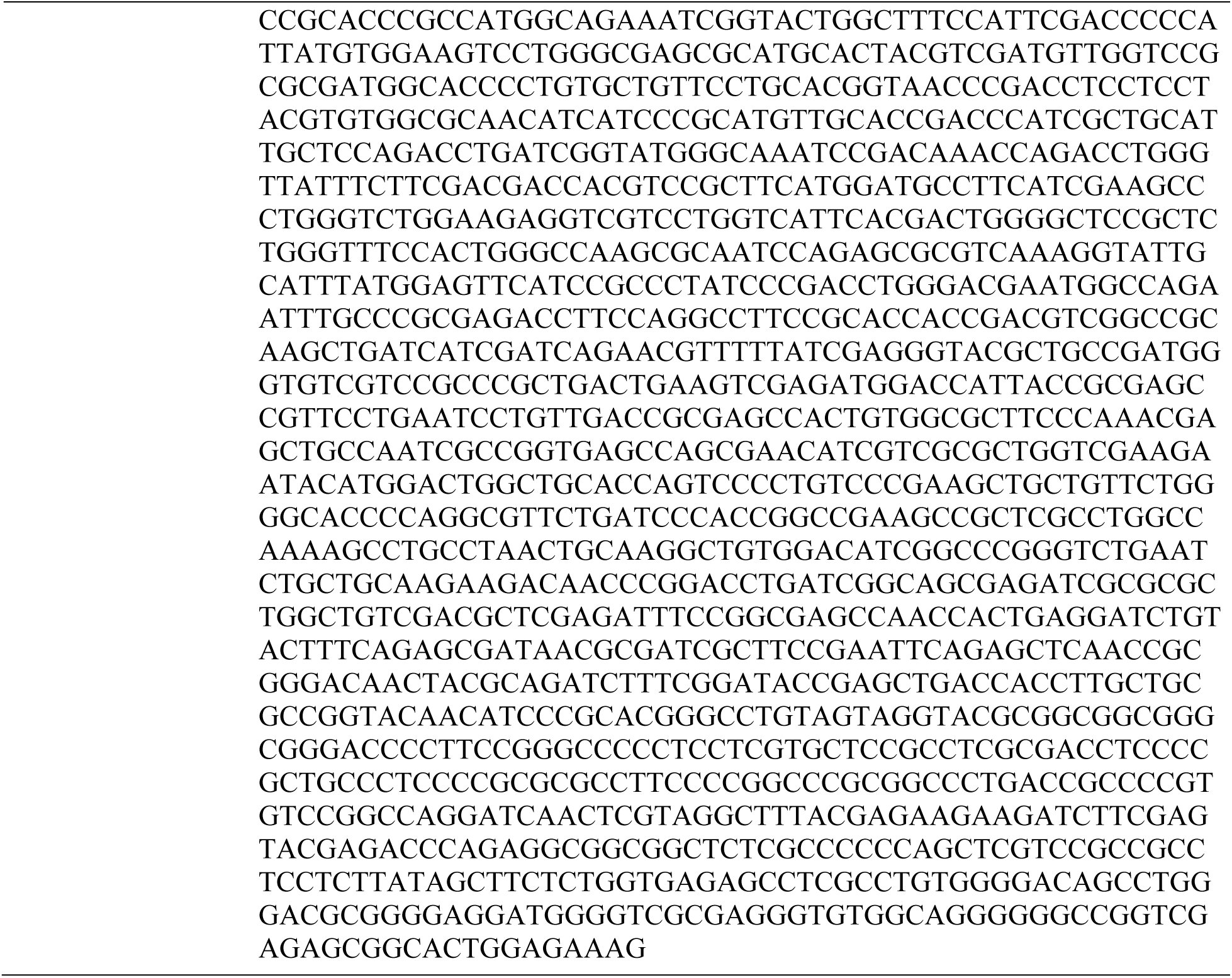
Homology-directed repair (HDR) template sequences used for the generation of stable cell lines used in this study.

**Table S3:**
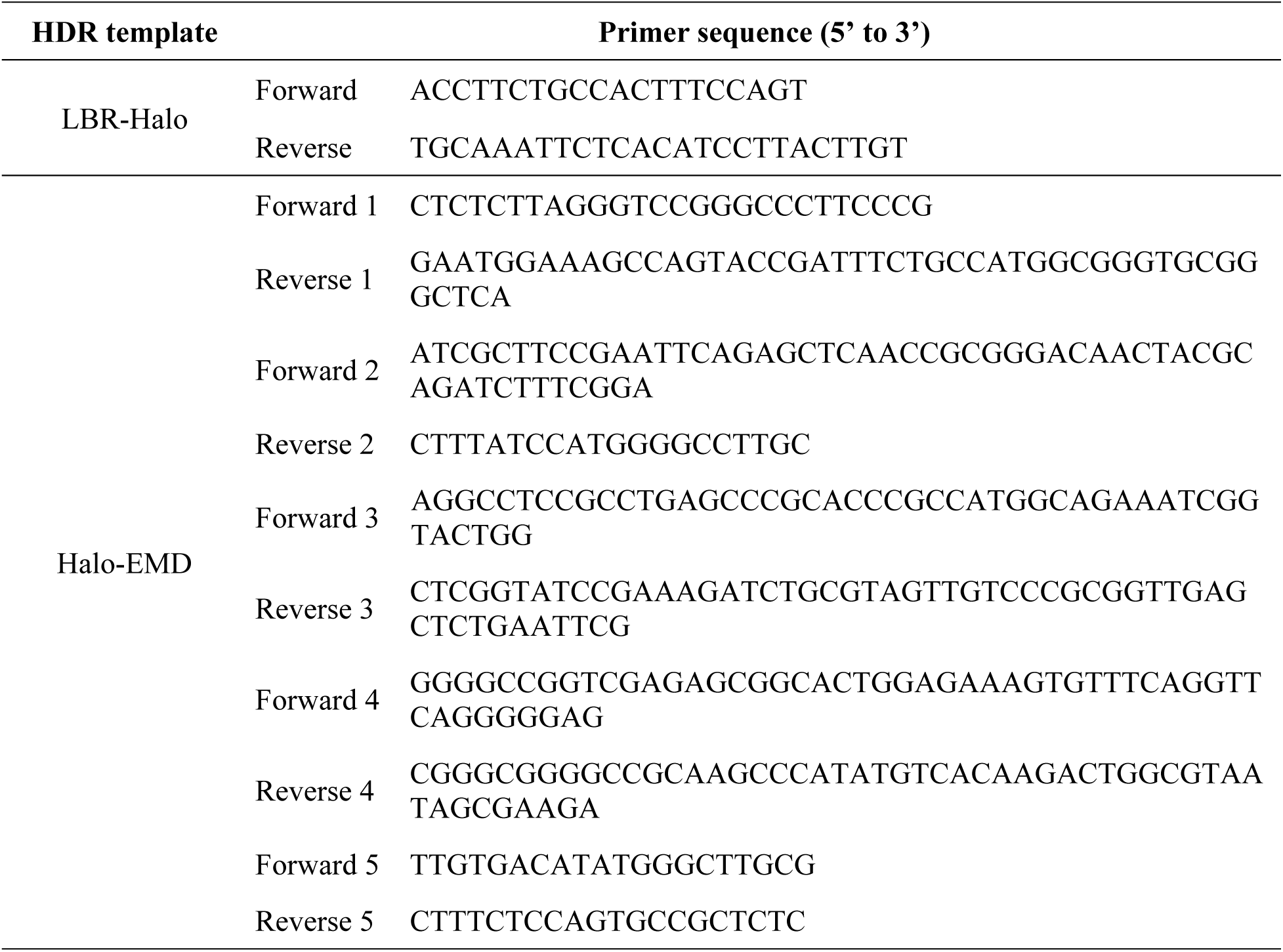
List of primers used for the generation of HDR templates used in this study.

